# Integrated rational and evolutionary engineering of genome-reduced *Pseudomonas putida* strains empowers synthetic formate assimilation

**DOI:** 10.1101/2022.07.10.499488

**Authors:** Justine Turlin, Beau Dronsella, Alberto De Maria, Steffen N. Lindner, Pablo I. Nikel

## Abstract

Formate is a promising, water-soluble C1 feedstock for biotechnology since it can be efficiently produced from CO_2_—but very few industrially-relevant hosts have been engineered towards formatotrophy. Here, the non-pathogenic soil bacterium *Pseudomonas putida* was adopted as a platform for synthetic formate assimilation. The metabolism of genome-reduced variants of *P. putida* was rewired to establish synthetic auxotrophies that could be functionally complemented by expressing components of the reductive glycine (rGly) pathway. The rGly pathway mediates the formate → glycine → serine transformations that yield pyruvate, ultimately assimilated into biomass. We adopted a modular engineering approach, dividing C1 assimilation in segments composed of both heterologous activities (sourced from *Methylorubrum extorquens*) and native reactions. Promoter engineering of chromosomally-encoded functions coupled to modular expression of rGly pathway elements enabled growth on formate as carbon source and acetate for energy supply. Adaptive laboratory evolution of two lineages of engineered *P*. *putida* formatotrophs significantly reduced doubling times to ca. 15 h. During evolution, two catabolic regimes became predominant in independently evolved clones, either *via* glycine hydroxymethylation (GlyA) or oxidation (ThiO). Taken together, our results expand the landscape of microbial platforms for C1-based biotechnological production towards supporting a formate bioeconomy.

**Graphical Abstract:** 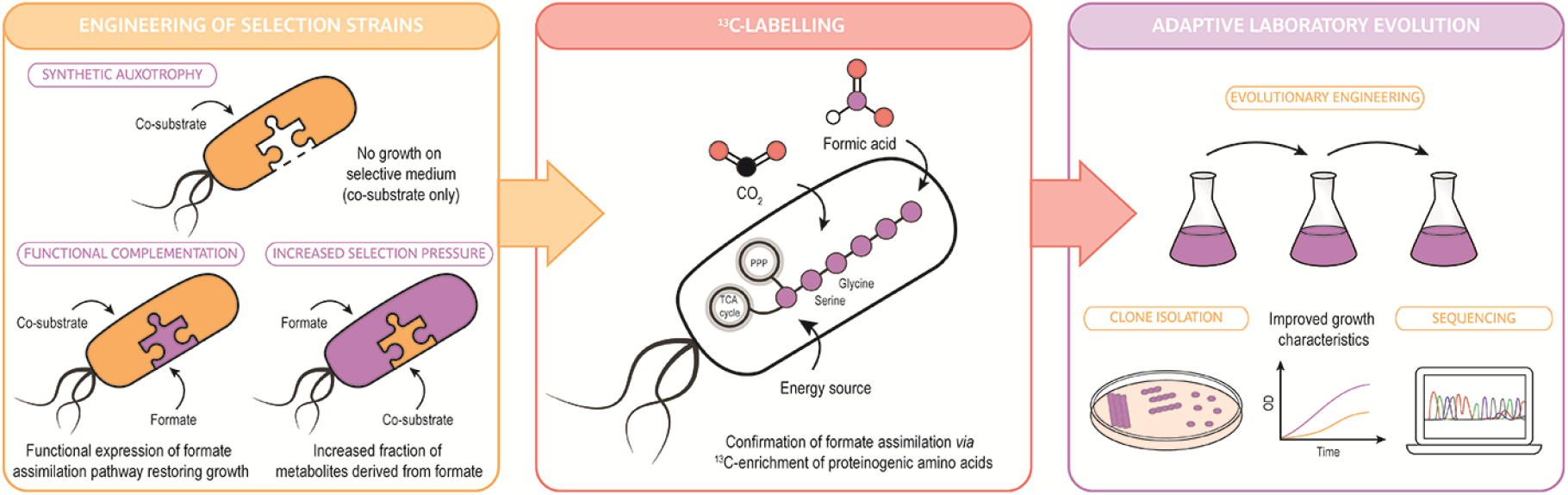

## 1. Introduction

Microbial synthesis of added-value fuels and chemicals continues to gather strength as an alternative to oil-based production—yet several drawbacks stand on the way towards widespread implementation and adoption of bioprocesses, e.g. limited supply of sugar-based feedstock, which competes with human consumption (1). Crop residues constitute a promising substitute of sugar substrates but, as of today, the pre-treatment and enzyme digestion steps needed for their use preclude the development of cost-efficient bioprocesses. In addition, extensive optimization is still required to improve yields on lignocellulose-derived fermentable sugars and to overcome the impact of inhibitors (2,3). CO_2_ and other C1 substrates, in contrast, hold the potential to replace fossil carbon as a feedstock to support bioproduction (4,5).

CO_2_ can be converted into added-value compounds by chemical processes that often involve harsh conditions, e.g. high temperature and pressure. In the biological realm, plants, algae and some bacteria are naturally able to fix CO_2_ *via* the (catalytically inefficient) Calvin-Benson-Bassham cycle and can be further engineered for the production of compounds of interest (6,7). The very nature of some gaseous substrates, however, imposes practical difficulties towards the implementation of production processes at a scale (5). From a biotechnological point of view, formic acid (HCOOH^1^) is an interesting C1 feedstock as it is highly soluble in H_2_O, which prevents gas-to-liquid mass transfer limitations. Besides, CO_2_ can be reduced electrochemically to formate with an energetic efficiency ≥40 (8–10). Alas, a very restricted number of microorganisms are able to grow on formate as the carbon source. Anaerobic acetogens, for instance, assimilate this compound *via* the Wood-Ljungdahl pathway (11,12). This highly energy-demanding route supports the production of a restricted range of small molecules mostly derived from acetyl-coenzyme A (CoA), since it can only yield compounds tied to ATP generation (13). The spectrum of C1-derived products is broader in the presence of O_2_ as generation of cellular energy and biosynthesis are uncoupled—which often comes at the price of a relatively low bioconversion efficiency (14). To tackle this issue, the linear reductive glycine (rGly) pathway has been identified as the most energy-efficient aerobic pathway supporting formate assimilation (7,15,16). In this pathway, formate is firstly condensed with tetrahydrofolate (THF) and reduced to methylene-THF.

The glycine cleavage system (GCS) then condenses the C1-moiety of methylene-THF with CO_2_ (thus capturing yet another C1 unit) and NH_3_ to yield glycine, completing the core reactions of the pathway. Next, glycine reacts with the C1-moiety of methylene-THF to form serine, subsequently deaminated to pyruvate (thereby feeding central carbon metabolism). The synthetic rGly route has been successfully implemented in two model bacterial species, *Escherichia coli* (17–20) and *Cupriavidus necator* (21), with the ultimate goal of building a formate bioeconomy (22). These studies illustrated the feasibility of incorporating synthetic C1 assimilation pathways in surrogate microbial hosts and pointed to some shortcomings related to the use of formate as a substrate (e.g. toxicity of the feedstock).

Owing to its rich and versatile metabolism, the Gram-negative, non-pathogenic soil bacterium *Pseudomonas putida* is a robust platform for bioproduction, capable of resisting oxidative stress and toxic chemicals (23–29). *P*. *putida* is known for its remarkable ability of processing and assimilating a wide variety of structurally unrelated substrates, ranging from simple sugars and organic acids to aromatic compounds and difficult-to-degrade xenobiotics (30)—yet, C1 compounds are excluded from its natural substrate ‘palate’ (31–33). Expanding the substrate range of *P*. *putida* to accommodate C1 feedstock by establishing a synthetic formatotrophic metabolism would enormously multiply the biotechnological value of this species. On this background, here we adopted a combined engineering approach to bestow formate assimilation *via* the synthetic rGly pathway onto reduced-genome strains of *P. putida*. Our strategy is rooted on creating synthetic auxotrophies that can be relieved by the functional expression of orthogonal metabolic modules and includes promoter engineering on native elements. Isotopic enrichment of proteinogenic amino acids in ^13^C-tracer experiments confirmed a formatotrophy pattern of C1 assimilation, and adaptive laboratory evolution (ALE) enabled efficient growth on the engineered *P*. *putida* strain using formate as the main carbon source. Hence, this work lays the ground for producing value-added compounds from soluble C1 substrates in *P*. *putida*—and contributes to establishing a true formate bioeconomy where diverse microbial platforms can be chosen for specific biotechnological purposes.

## 2. Materials and Methods

### 2.1. Chemicals and reagents

Chemicals were purchased from Sigma-Aldrich Co. (St. Louis, MO, USA) unless otherwise indicated. Oligonucleotides were synthesized by Integrated DNA Technologies Inc. (Coralville, IA, USA) and PCR reactions were performed using Phusion^TM^ or Phusion *U* Hot Start^TM^ DNA polymerases from ThermoFisher Scientific Co. (Waltham, MA, USA). PCR fragments were cloned into plasmids with either the USER^TM^ enzyme (ThermoFisher Scientific Co.), T4 polynucleotide kinase or T4 DNA ligase from New England BioLabs (Ipswich, MA, USA). DNA sequencing was performed at Eurofins Genomics (Ebersberg, Germany). All primers used in this study are listed in **Table S1** in the Supplemental Data.

### 2.2. Bacterial strains and medium composition

All bacterial strains and plasmids are listed in **Tables 1** and **2**, respectively. *E. coli* DH5α and *E. coli* DH5α λ*pir* were used as cloning hosts, and *P*. *putida* KT2440 and its reduced-genome derivative EM42 were used for engineering purposes. Lysogeny broth (LB) complex medium (containing 10 g L^−1^ tryptone, 5 g L^−1^ yeast extract and 10 g L^−1^ NaCl) and de Bont minimal (DBM) medium were used for strain cultivation (34). DBM medium contained 3.88 g L^−1^ K_2_HPO_4_, 1.63 g L^−1^ NaH_2_PO_4_, 2 g L^−1^ (NH_4_)_2_SO_4_, 0.1 g L^−1^ MgCl_2_·6H_2_O, with initial pH adjusted at 7.0. DBM medium was added with a trace elements solution [10 mg L^−1^ ethylenediaminetetraacetic acid (EDTA), 2 mg L^−1^ ZnSO_4_·7H_2_O, 1 mg L^−1^ CaCl_2_·2H_2_O, 5 mg L^−1^ FeSO_4_·7H_2_O, 0.2 mg L^−1^ Na_2_MoO_4_·2H_2_O, 0.2 mg L^−1^ CuSO_4_·5H_2_O, 0.4 mg L^−1^ CoCl_2_·6H_2_O, 1 mg L^−1^ MnCl_2_·2H_2_O] and the carbon sources indicated in the text. Solid culture media additionally contained 15 g L^−1^ agar. When needed, kanamycin (Km), gentamicin (Gm) and streptomycin (Str) were added at 50 μg mL^−1^, 10 μg mL^−1^ and 100 μg mL^−1^, respectively.

**Table 1.** Bacterial strains used in this study.

| Strain | Relevant characteristics<sup>a</sup> | Reference or source |
| --- | --- | --- |
| *Escherichia coli* |  |  |
| DH5α | Cloning host; F<sup>-</sup> λ<sup>-</sup> *endA1 glnX44(AS) thiE1 recA1 relA1 spoT1 gyrA96(Nal<sup>R</sup>) rfbC1 deoR nupG Φ80(lacZΔM15) Δ(argF-lac)U169 hsdR17(r<sub>K</sub><sup>-</sup> m<sub>K</sub><sup>+</sup>)* | Hanahan and Meselson (69) |
| DH5αλ<sub>pir</sub> | Cloning host; F<sup>-</sup> λ<sup>-</sup> *endA1 glnX44(AS) thiE1 recA1 relA1 spoT1 gyrA96(Nal<sup>R</sup>) rfbC1 deoR nupG Φ80(lacZΔM15) Δ(argF-lac)U169 hsdR17(r<sub>K</sub><sup>-</sup> m<sub>K</sub><sup>+</sup>)*, λ<sub>pir</sub> lysogen | Platt et al. (70) |
| MG1655 | Wild-type strain, derivative of K-12, F<sup>-</sup> λ<sup>-</sup> *ilvG rfb-50 rph-1* | Blattner et al. (71) |
| *Pseudomonas putida* |  |  |
| KT2440 | Wild-type strain, derived from *P. putida* mt-2 (72) cured of the TOL plasmid pWW0 | Bagdasarian et al. (73) |
| EM42 | Reduced-genome derivative of *P. putida* KT2440; Δ*PP_4329-PP_4397* (flagellar operon) Δ*PP_3849-PP_3920* (prophage I) Δ*PP_3026-PP_3066* (prophage II) Δ*PP_2266-PP_2297* (prophage III) Δ*PP_1532-PP_1586* (prophage IV) Δ*Tn7* Δ*endA-1* Δ*endA-2* Δ*hsdRMS* Δ*Tn4652* | Martínez-García et al. (53) |
| SL | Derivative of *P. putida* EM42, Δ*serA* Δ*ltaE* | This study |
| SLA | Derivative of *P. putida* SL, Δ*aceA* | This study |
| AceA | Derivative of *P. putida* EM42, Δ*aceA* |  |
| pM1 | Derivative of *P. putida* SLA carrying plasmid pSFM1 (P<sub>4</sub>→*mtdA-fch-fftL* from *Methylobacterium extorquens*); Km<sup>R</sup> | This study |
| pM1gM2 | Derivative of *P. putida* SLA, Δ*PP_0997* Δ*PP_5002-PP_5008* P<sub>14g</sub>(*BCD10*)→*gcv-I* carrying plasmid pSFM1; Km<sup>R</sup> | This study |
| gM1gM2 | Derivative of *P. putida* SLA, Δ*PP_0997* P<sub>14g</sub>(*BCD10*)→*gcv-I* Δ*PP_5002-PP_5008*::P<sub>4</sub>→*mtdA-fch-fftL* from *M. extorquens* | This study |
| rG1 | Derivative of *P. putida* gM1gM2, $P_{14g}(BCD2) \rightarrow glyA-I$<br>$P_{14g}(BCD2) \rightarrow glyA-II$ | This study |
| rG2 | Derivative of *P. putida* gM1gM2, $\Delta crc$ $\Delta hexR$ $\Delta purT$<br>$P_{14g}(BCD10) \rightarrow gcv-II$ $P_{EM7} \rightarrow pntAA$ $P_{14g}(BCD20) \rightarrow glyA-I$<br>$P_{14g}(BCD20) \rightarrow glyA-II$ carrying plasmid pSFM3; Km<sup>R</sup> | This study |
| rG1_1 | Evolved population of *P. putida* rG1 | This study |
| rG1_2 | Evolved population of *P. putida* rG1 | This study |
| rG2_1 | Evolved population of *P. putida* rG2 | This study |
| rG2_2 | Evolved population of *P. putida* rG2 | This study |
| rG1_1/2 | Isolated clone from the evolved *P. putida* rG1_1 population | This study |
| rG2_2/5 | Isolated clone from the evolved *P. putida* rG2_2 population | This study |
| rG1_1/2 $\Delta thiO$ | Derivative of *P. putida* rG1_1/2, $\Delta thiO$ | This study |
| rG2_2/5 $\Delta thiO$ | Derivative of *P. putida* rG2_2/5, $\Delta thiO$ | This study |

a Antibiotic markers: Km, kanamycin; and Nal, nalidixic acid.

### 2.3. Construction of plasmids and engineered strains

#### 2.3.1. P. putida gene deletions and genome integrations

General methods for genome and gene manipulations followed well established protocols (35–39). Gene deletions in *P. putida* EM42 were performed by I-*Sce*I–mediated recombination using flanking sequences amplified from the genome of strain EM42 and cloned into the suicide pGNW2 plasmid (40–42). To this end, 500-bp DNA fragments immediately upstream and downstream of the locus to be deleted were amplified from *P. putida* EM42 with Phusion *U* Hot Start^TM^ DNA polymerase using uracil-containing primers. Plasmid pGNW2 was amplified *via* the same method and digested with *Dpn*I. Next, 100 ng of each PCR fragment were mixed with 1 μL of *Dpn*I-treated pGNW2 plasmid and 1 μL of USER^TM^ enzyme in a final volume of 10 μL. The reaction was incubated for 30 min at 37°C, followed by a decrease in the temperature during 3 min (from 28°C to 20°C, 2°C per step), and an incubation at 10°C for at least 10 min. A 5-μL aliquot of the USER mix was transformed into chemically-competent *E. coli* DH5a l*pir* and, upon recovery, the cell suspension was plated onto LB medium agar plates containing the corresponding antibiotic. These operations were used to construct suicide plasmids pGNW2·Δ*serA*, pGNW2·Δ*ltaE*, pGNW2·Δ*aceA*, pGNW2·Δ*PP_0997,* pGNW2·Δ*PP_1236,* pGNW2·Δ*crc,* pGNW2·Δ*purT,* pGNW2·Δ*gcv-I,* pGNW2·Δ*gcv-II,* pGNW2·Δ*glyA-I,* pGNW2·Δ*glyA-II,* pGNW2·Δ*thiO,* pGNW2·Δ*tdcG-I,* pGNW2·Δ*tdcG-II* and pGNW2·Δ*tdcG-III* (**Table 2**). Positive clones were identified by colony PCR and verified by DNA sequencing.

**Table 2.**
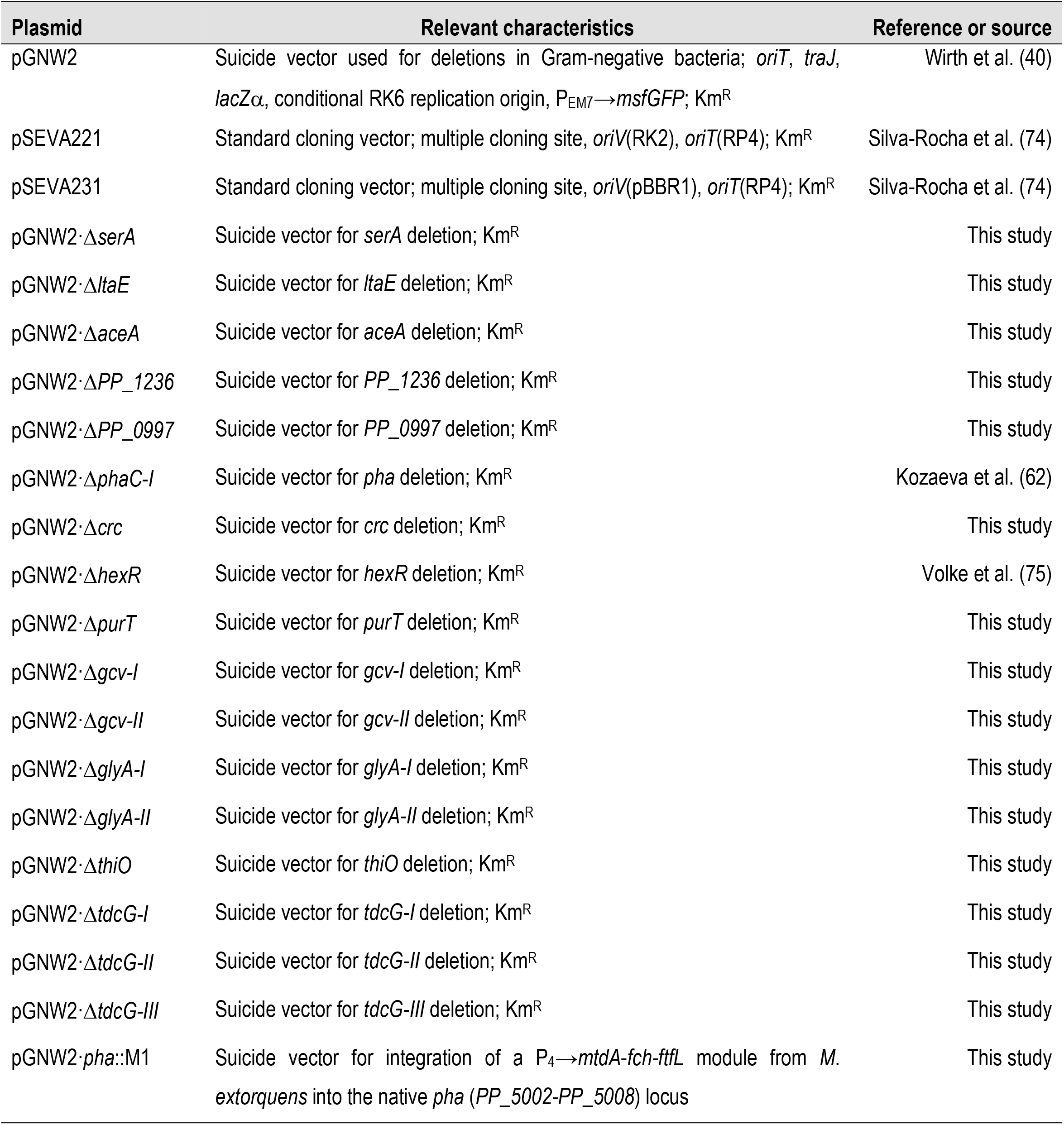
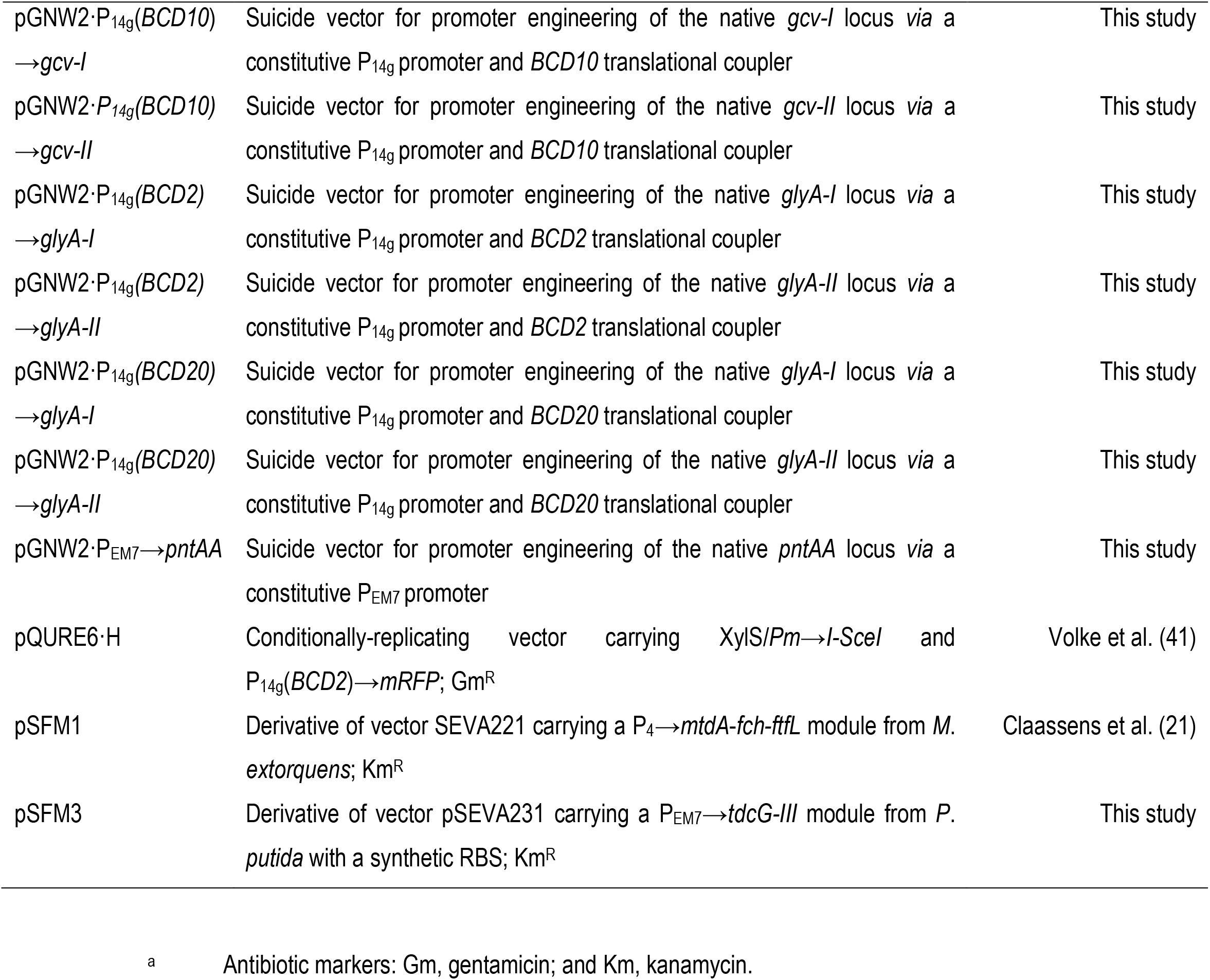
Plasmids used in this study.

The suicide pGNW2-derivative plasmids, bearing homologous regions needed for chromosomal integration, were delivered into the cells by electroporating 50 μL of freshly-prepared *P. putida* EM42 electrocompetent cells (washed three times with 300 mM sucrose) with 300 ng of DNA. Electroporation was performed with a Gene Pulser XCell (Bio-Rad, Hercules, CA, USA) set to 2.5 kV, 25 μF capacitance and 200 Ω resistance in a 2-mm gap cuvette. After selecting positive co-integration events, a conditionally-replicative plasmid bearing the meganuclease gene I-*Sce*I was introduced into the cells *via* electroporation. The I-*Sce*I meganuclease led to restriction of pGNW2 co-integrants within the chromosome and forced a second homologous recombination event. Cells were recovered in 1 mL of LB medium and 2 mM of 3-methylbenzoate (3-*m*Bz) for at least 3 h at 30°C, and plated onto LB medium agar containing the corresponding antibiotic(s) and 1 mM 3-*m*Bz to induce plasmid replication and I-*Sce*I expression (43). Positive clones were identified by colony PCR, verified by DNA sequencing and cured from the resolving plasmid by serial dilution under non-selective conditions.

#### 2.3.2. Genome integration of ftfL, fch and mtdA in a knock-in/knock-out strategy

Plasmid pSFM1 contains the *ftfL*, *fch* and *mtdA* genes from *M. extorquens* AM1 preceded by synthetic RBS under the control of the P_4_ promoter in a pSEVA221 vector (**Table 2**). Plasmid pSFM1 was used as a template to obtain codon-optimized gene sequences used for engineering *P*. *putida* as indicated in **Table S2** in the Supplemental Material. For genome-based expression of *ftfL*, *fch* and *mtdA*, the synthetic operon was inserted into the *pha* locus of *P*. *putida* by amplification with Phusion *U* Hot Start^TM^ DNA polymerase using uracil-containing primers followed by USER cloning yielding pGNW2·*pha*::M1. Chromosomal integration of this segment was performed as described in the section above.

#### 2.3.3. Promoter engineering of chromosomally-encoded genes and synthetic modules

Genomic overexpression of the *gcv-I/II* operons, *glyA-I/II* and *pntAB* was achieved using the previously described I-*Sce*I–mediated recombination. The integration plasmids for promoter engineering were constructed by amplifying the first 500-bp of the locus, the 500-bp upstream of the operon and the synthetic promoter or translational coupler (as a bicistronic design, BCD) and inserting the amplicons in the pGNW2 suicide plasmid by USER cloning (44). This operation yielded pGNW2·P_14g_(*BCD2*)→*glyA-I* and pGNW2·P*_14g_(BCD2)*→*glyA-II* (**Table 2**). Plasmids for promoter engineering of pGNW2·P_14g_(*BCD20*)→*glyA-I*, pGNW2·P_14g_(*BCD20*)→*glyA-II*, *gcv-I*, *gcv-II*, *pntAB*, and *tdcG-III* were constructed by replacing regulatory sequences in previously constructed plasmids by Quick Change (45), amplifying the entire plasmid with forward and reverse primers containing the new promoter sequence (and BCDs, when relevant). The amplified product was restricted with *Dpn*I, phosphorylated and ligated. Phosphorylation and ligation were done using T4 polynucleotide kinase and T4 DNA ligase, respectively, according to the manufacturer’s protocol, giving rise to plasmids pSFM3, pGNW2·P_14g_(*BCD20*)→*glyA-I*, pGNW2·P_14g_(*BCD20*)→*glyA-II*, pGNW2·P_14g_(*BCD10*)→*gcv-I*, pGNW2·P_14g_(*BCD10*)→*gcv-II* and pGNW2·P_EM7_-*pntAA*. The plasmids were transformed into *E. coli* DH5a l*pir* and cells were plated onto LB medium agar containing the corresponding antibiotic. Positive clones were identified by colony PCR and verified by DNA sequencing before transferring the integration vectors into *P*. *putida* as described above.

### 2.4. Growth conditions for C1 assimilation experiments

Overnight cultures of the engineered strains in LB medium were diluted 100× times to inoculate a preculture in 2-5 mL of DBM medium supplemented with 20 mM glucose and 2 mM glycine or 30 mM formate and 10% (v/v) CO_2_ in the headspace in a 12-mL culture tube and incubated at 30°C at 250 rpm. The overnight culture was collected at 12,000×*g* for 2 min and washed twice with DBM medium to inoculate the main culture in either 12-mL culture tubes, 96-well microtiter plates or baffled shaken-flasks with the appropriate carbon source(s). For growth in 96-well microtiter plates, 200 μL of a cell suspension at an optical density measured at 600 nm (OD_600_) of 0.05 were incubated with orbital shaking in a Synergy H1 microtiter plate reader (BioTek Instruments Inc., Winooski, VT, USA) or in an ELx808 microtiter plate reader (BioTek Instruments Inc.). Cultivations in test tubes were conducted in 2.5 mL of medium at 30°C and 200 rpm, and shaken-flask cultivations were done in 30 mL of medium in a 250-mL Erlenmeyer flask at 30°C and 150 rpm. Depending on the experiment, DBM medium was added with 20 mM glucose, 30 or 60 mM formate, 24 mM ribose and/or 20 mM acetate, and, whenever relevant, cultivations were done in a CO_2_-enriched atmosphere. To this end, 10% (v/v) CO_2_ was flushed in the headspace by built-in gas controllers. The specific growth rate (μ) was estimated during exponential growth by linear regression on the average OD_600_ values and doubling times (DTs, in h) were calculated according to DT = *ln*(2)/μ.

### 2.5. Formate toxicity assays

Formate toxicity was assessed by drop assays with overnight cultures of strains *P. putida* KT2440, EM42 and *E. coli* MG1655 grown in LB medium. Cells were washed with DBM medium without carbon source and serially diluted in fresh DBM medium. Finally, 20 μL of each dilution was spotted onto LB agar plates supplemented with 0, 60, 120, 180 or 240 mM formate and incubated for 24 h at 30°C for *P. putida* or 37°C for *E. coli*. Survival percentages correspond to the ratio of the number of colony forming units (CFUs) in the presence of formate normalized by the number of CFUs without formate at the 10^-6^ dilution. The washed overnight cultures of *P. putida* KT2440, EM42 and *E. coli* MG1655 were also incubated in 96-well plates (flat bottom; Greiner Bio-One, Kremsmünster, Austria) in 200 μL of LB medium inoculated at OD_600_ 0.05 in an ELx808 microtiter plate reader (BioTek Instruments Inc.) with 0, 120 or 180 mM formate at 30 or 37°C to assess toxicity in liquid medium.

### 2.6. Adaptive laboratory evolution (ALE) experiments

Overnight cultures in LB medium were diluted 100× to inoculate a preculture in 2-5 mL of DBM medium supplemented with 20 mM glucose, 30 mM formate and 10% (v/v) CO_2_ in the headspace. The washed preculture was used to inoculate the main culture at an OD_600_ of 0.05 in 2-5 mL of DBM medium supplemented with 30 mM formate, 20 mM acetate and 10% (v/v) CO_2_ in the headspace in a 12-mL culture tube and incubated at 30°C at 200 rpm. The culture was passaged serially in a new 12-mL culture tube containing fresh medium (with the same additives indicated above) until reaching stationary phase. Isolates rG1_1 and rG1_2, as well as rG2_1 and rG2_2, indicate bacteria evolved in parallel ALE experiments with the same strain. After 5 and 6 passages under formatotrophic conditions, evolved populations of *P*. *putida* rG1 and rG2, respectively, were streaked onto DBM medium plates supplemented with 30 mM formate and 20 mM acetate and incubated in the presence of 10% (v/v) CO_2_ in the headspace in order to isolate individual clones. Clones were further screened for reduced doubling time in 96-well microtiter plates.

### 2.7. Whole-genome sequencing

Genomic DNA was extracted from *P*. *putida* from 2 mL of overnight LB medium cultures with the PureLink Genomic DNA Mini kit (ThermoFisher Scientific Co.). Library construction, sequencing, and subsequent data quality control were performed by Novogene Co. Ltd. (Cambridge, United Kingdom). Sequencing was performed using the Illumina NovaSeq 6000 PE150 platform to obtain 150 bp paired-end reads. Reads were paired, trimmed with the *BBDuk* plugin, normalized and assembled to *P. putida* EM42 reference genome using the Geneious 2021.1.1 software (Dotmatics, Boston, MA, USA). The trimming procedure included (i) adapters using presets for Illumina adapters, (ii) read ends based on quality (Q>20), (iii) adapters based on paired read overhangs and (iv) short reads (<20 bp). Mutations with a frequency above 50% and excluded from low coverage regions were identified after comparison with the pre-evolved strains as described previously (46,47).

### 2.8. Analytical procedures

Extracellular formate, glucose and acetate concentrations were measured in culture supernatants after centrifuging the cultures for 5 min at 10,000×*g*. Supernatants were analyzed in a high-performance liquid chromatography instrument (UltiMate™ 3000 Basic Automated System, ThermoFisher Scientific Co.) equipped with an Aminex HPX-87P column (Bio-Rad) and a refractive index detector Shodex RI-101 (Showa Denko America Inc., NY, USA). Specific glucose (*q*_G_) and formate (*q*_F_) consumption rates were determined with the following equation:

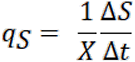

where *q_S_* is the biomass-specific substrate consumption rate (mmol · g_CDW_^−1^ · h^−1^), *X* is the average biomass concentration between two sampling time points (g_CDW_ · L^−1^), Δ*S* is the difference in substrate concentration between two sampling time points (mM) and Δ*t* is the time between two sampling points (h). The *q*_G_ value corresponds to the growth rate divided by the slope of the linear regression of the biomass concentration over glucose concentration. The *q*_G_ value reported is an average of the values determined individually for three biological replicates; the specific formate consumption rate was estimated for each timepoint by a spline fit in the independent experiments.

### 2.9. ^13^C-isotopic labeling of proteinogenic amino acids

A preculture was inoculated in 5 mL LB medium and incubated for 24 h at 30°C. The next day, the cells were inoculated in DBM medium with the appropriate carbon source (20 mM glucose, 24 mM ribose, 20 mM acetate and 30 or 60 mM formate) and incubated with 10% (v/v) of CO_2_ in the headspace. The precultures were washed three times with DBM medium without carbon source and inoculated in the same medium but replacing the non-labeled carbon source with ^13^C-sodium formate (Sigma-Aldrich Co.), 1-^13^C-sodium acetate (Cambridge Isotope Laboratories Inc., Andover, MA, USA) and/or ^13^CO_2_ (Cambridge Isotope Laboratories Inc.) when indicated. Labeling experiments were performed in 24-well deep-well plates with a working volume of 2.5 mL and at an initial OD_600_ of 0.03. For cultures with ^13^CO_2_, experiments were performed in 3 mL in tubes at an initial OD_600_ of 0.01 in an airtight 6 L-desiccator (Lab Companion, Daejeon, Korea). The desiccator was flushed with 10% (v/v) ^13^CO_2_ and 90% air on a shaker platform (180 rpm). Once the cells reached stationary phase, an equivalent of 1 mL of culture at an OD_600_ of 0.8 were harvested and washed once with MilliQ H_2_O and hydrolyzed in 1 mL of 6 M HCl for 24 h at 95°C (48) HCl was removed *via* heating at 95°C for another 12 h with open lids. The hydrolyzed biomass was then resuspended in 1 mL of double-deionized H_2_O. Ultra-performance liquid chromatography (Acquity, Milford, MA, USA) with a C_18_-reversed-phase column (Waters, Eschborn, Germany) was used to separate amino acids in the hydrolysate. Mass spectra were obtained using an Exactive mass spectrometer (ThermoFisher Scientific Co.), and data was examined using the Xcalibur software (ThermoFisher Scientific Co.). Amino acid standards (Merck, Darmstadt, Germany) were analyzed beforehand to determine typical retention times.

### 2.10. Statistical analysis

Data analysis was performed with MS Excel and Prism 8 (GraphPad Software Inc., San Diego, CA, USA) unless differently specified. Reported values are indicated as averages ± standard deviation of replicates as specified in the legend to the corresponding figures. When relevant, the level of significance of differences when comparing results was evaluated by ANOVA (Dunnett’s test, Prism 8) with α < 0.05.

## 3. Results and Discussion

### 3.1. Pseudomonas putida as a robust host for engineering synthetic formate assimilation

Efficient bioprocesses based on formate as a feedstock call for the use of high concentrations of the C1 substrate to promote sufficient bacterial growth and biosynthesis of the target product(s) (49). This requirement is challenged by the fact that formate hampers microbial growth due to the diffusion of the protonated form [pK*_a_* (HCOOH) = 3.74] across the cell membrane, acidifying the cytoplasm and negatively affecting the H^+^ motive force (50). Formate also imposes oxidative stress on Gram-negative bacteria (51). These observations prompted the question of whether *P*. *putida* can tolerate formate at concentrations compatible with its use as a carbon substrate. To address this point, bacterial cell suspensions were spotted on (rich) LB medium plates containing different formate concentrations up to 240 mM (**Fig. 1**). Wild-type *P*. *putida* KT2440 thrived on LB medium even at the highest concentration tested (**Fig. 1a**), with only a minor decrease in the colony forming units (CFUs) count when compared to the control experiment (i.e. no added formate). Since *E. coli* has been recently engineered towards synthetic formatotrophy (17,18), this species (wild-type strain MG1655) was also included in these experiments as a comparison. The growth of *E*. *coli* on LB medium plates was severely reduced by formate at 180 mM, and completely abolished at 240 mM (**Fig. 1a**). These findings were quantitatively confirmed by cultivating the two species in liquid LB medium with formate (**Fig. 1b**), which indicated that >70% of the *P*. *putida* population can survive a 180 mM challenge. The impact of the C1 substrate on bacterial growth was further evidenced by a concentration-dependent decrease in the doubling time (DT) of *E*. *coli* MG1655 (**Fig. 1c**), ca. 5-fold longer in the presence of 180 mM formate than in control experiments. The growth patterns of *P*. *putida* (both wild-type KT2440 and its reduced-genome derivative EM42), on the contrary, showed no significant differences in the presence of 120 or 180 mM formate as compared to control experiments (**Fig. 1d**). These results highlight the value of *P*. *putida* as a surrogate host of synthetic formatotrophy routes due to its high tolerance to stressful conditions (52)—in this case, brought about by the feedstock. Owing to its emergent phenotypic and metabolic properties [e.g. enhanced ATP and NADPH availability (53,54)], and since DTs were not affected by formate up to ca. 200 mM, we adopted strain EM42 as the *chassis* for engineering formate assimilation as explained in the next section.

**Figure 1.**
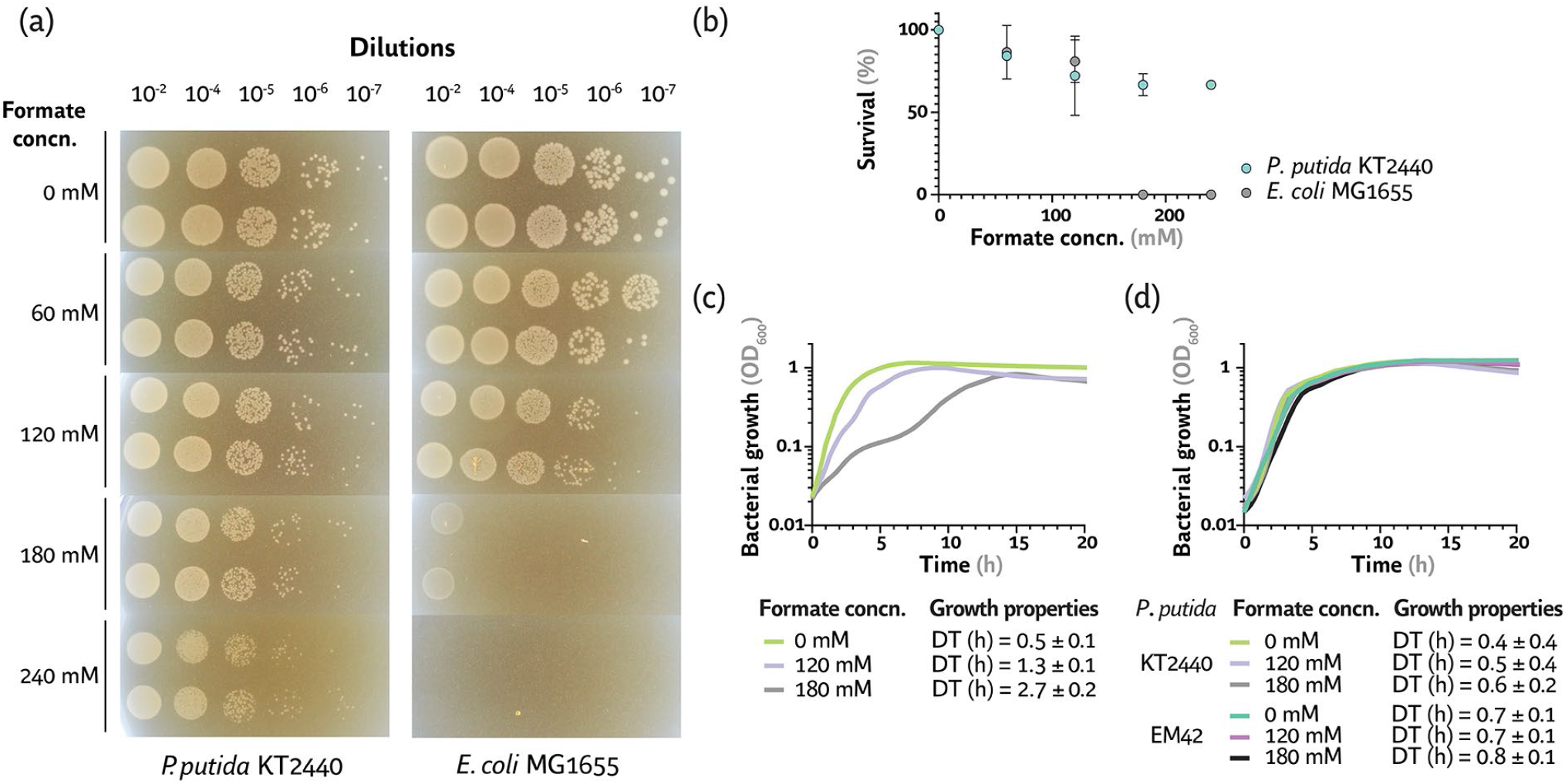
Evaluating formate toxicity in wild-type *E. coli* and *P. putida*. **(a)** Growth phenotypes of *E. coli* MG1655 and *P. putida* KT2440 exposed to increasing formate concentrations (concn.). Tolerance was assessed by spotting serially-diluted culture aliquots onto LB medium plates added with the C1 compound (as sodium salt) up to 240 mM. Plates were photographed after 24 h incubation either at 30°C (*P*. *putida*) or 37°C (*E*. *coli*). **(b)** Survival of the bacterial species under study upon exposure to formate for 24 h. Survival percentages were calculated as a fraction from the average of colony forming units (CFUs) in LB medium plates containing formate and normalized to the CFUs count in the control condition (i.e. no formate added) in two biological replicates. Growth profile of **(c)** *E. coli* MG1655 and **(d)** *P. putida* KT2440 and its reduced-genome derivative EM42 in liquid cultures containing formate. Cells were cultivated in microtiter plate cultures in LB medium supplemented with 0, 120 or 180 mM formate. Average values for bacterial growth (estimated as the optical density measured at 600 nm, OD_600_) and doubling time (DT) ± standard deviation of three biological replicates are represented in each case.

### 3.2. Establishing synthetic formate conversion in a serine auxotroph of P. putida

The general strategy adopted for establishing formate assimilation in *P*. *putida* is based on the creation of synthetic auxotrophies that can be relieved upon introducing specific pathway activities (55,56). Since the heterologous activities are directly coupled to bacterial growth, this approach offers the advantage of confirming the functionality of individual metabolic modules while identifying optimal expression levels. In this sense, the synthetic rGly pathway is divided in four functional metabolic modules (**Fig. 2**). The first module (M1, attachment of C1 to THF) converts formate into methylene-THF *via* the action of a set of three heterologous enzymes from *Methylobacterium extorquens* [i.e. formate-THF ligase (FtfL), methenyl-THF cyclohydrolase (Fch) and methylene-THF dehydrogenase (MtdA)] and yields 5,10-methylene-THF. The second module (M2, glycine generation) condenses 5,10-methylene-THF, NH_3_ and CO_2_ to form glycine by the native glycine cleavage system (GCS). In the third module (M3, pyruvate synthesis), the endogenous L-serine hydroxymethyltransferase (GlyA) further condenses glycine with yet another 5,10-methylene-THF unit to form serine, which is deaminated into pyruvate through the action of (native) L-serine deaminase (TdcG). Pyruvate then feeds central carbon metabolism through both gluconeogenesis and downward catabolism in the tricarboxylic acid (TCA) cycle.

**Figure 2.**
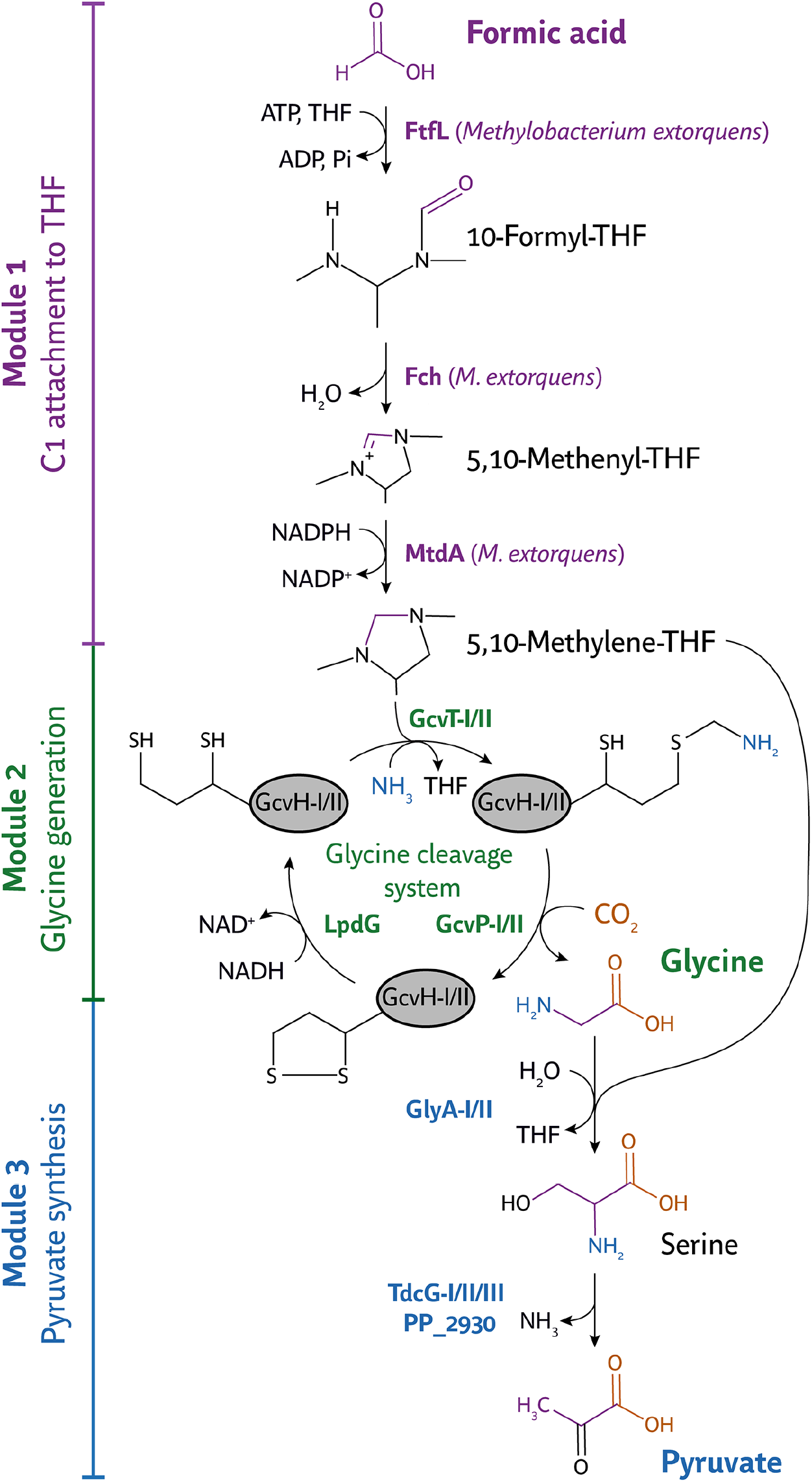
Biochemical structure of the synthetic reductive glycine (rGly) pathway. The route is divided in three modules, indicated on the left side of the scheme, composed by both endogenous and heterologous enzymatic activities. In the latter case, the source of the genes used in this study is indicated alongside the corresponding biochemical step. Note that two sets of genes encode GlyA, GcvT, GcvP and GcvH and three different genes encode TdcG in *P*. *putida* KT2440. For the sake of simplicity, a single substructure is represented for tetrahydrofolate (THF)-containing intermediates; the lipoic acid moiety of the H-protein in the native glycine cleavage system (GcvH) is explicitly shown. Key enzymatic activities in the pathway are indicated as follows: FtfL, formate-THF ligase; Fch, methenyl-THF cyclohydrolase; MtdA, methylene-THF dehydrogenase; GcvH, glycine cleavage system protein H; GcvP, glycine cleavage system protein P; GcvT, glycine cleavage system protein T; LpdG, dihydrolipoyl dehydrogenase; GlyA, serine hydroxymethyl transferase; and TdcG, L-serine deaminase.

The first step of our engineering strategy consisted in establishing the upper part of the synthetic rGly pathway (i.e. from formate to glycine) by establishing C1, glycine and serine auxotrophy in *P. putida* EM42 (**Fig. 3a**). Deletion of *serA* (*PP_5155*, encoding a bifunctional D-3-phosphoglycerate dehydrogenase/α-ketoglutarate reductase) and *ltaE* (*PP_0321*, encoding a low-specificity L-threonine aldolase) rendered strain EM42 unable to synthesize glycine or serine from glucose. In this glycine auxotroph (*P*. *putida* SL; **Table 1**), eliminating SerA prevents serine biosynthesis and removing LtaE blocks glycine synthesis from threonine. Furthermore, *aceA* (*PP_4116*, encoding isocitrate lyase, the first step of the glyoxylate shunt) was deleted to inhibit the potential formation of glycine from glyoxylate, leading to *P*. *putida* SLA. Growth phenotypes of this strain were tested in de Bont minimal medium (DBM) supplemented with 20 mM of glucose, in the absence or presence of amino acid additives (**Fig. S1** in the Supplemental Data). Strain SL (Δ*serA* Δ*ltaE*) was expectedly unable to grow after inoculation in DBM medium with glucose as the sole carbon source, but addition of glycine (**Fig. S1a**) or serine (**Fig. S1b**) restored growth in a concentration-dependent fashion. Supplementing glycine at 15 mM or serine at 7.5 mM was sufficient to reestablish a growth profile comparable to that of wild-type strain EM42. The glycine- and serine-dependence of strain SLA (Δ*serA* Δ*ltaE* Δ*aceA*) in DBM cultures was essentially the same as for *P*. *putida* SL (data not shown). Interestingly, *P*. *putida* can tolerate high serine concentrations (unlike *E*. *coli*), underlying the value of the host in this study for engineering C1 assimilation via this amino acid.

**Figure 3.**
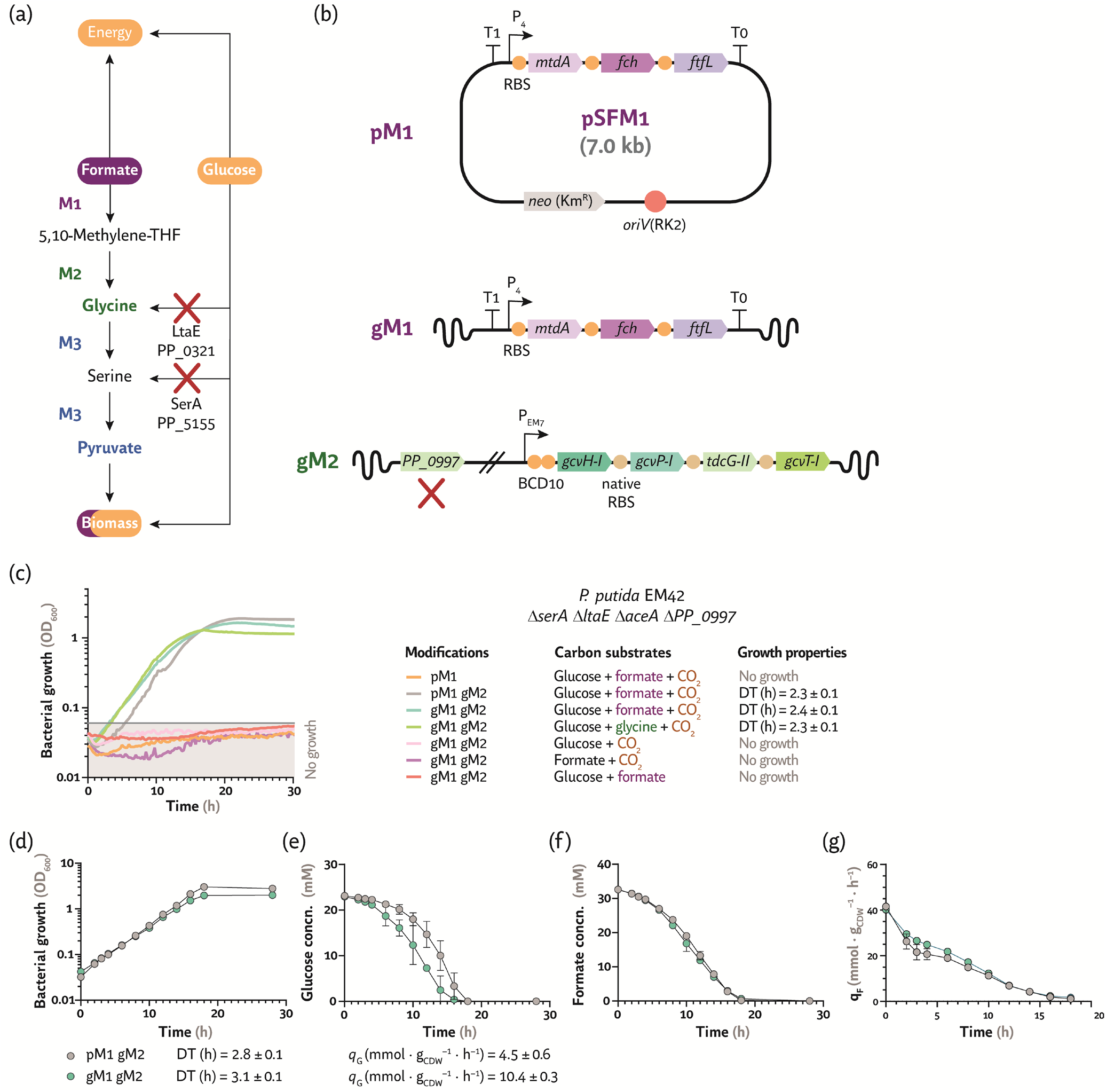
Engineering *P*. *putida* for formate assimilation in the presence of glucose and CO2. **(a)** Selection strategy to force formate assimilation into glycine and serine by deleting *ltaE* and *serA* (*P*. *putida* SL) and overexpressing modules M1 and M2. **(b)** Key elements implemented for gene and genome engineering of *P*. *putida* towards formatotrophy. Synthetic module M1 was expressed either from a low-copy-number, broad-host-range plasmid (pSFM1, **Table 1**) or directly from the genome by a knock-in/knock-out strategy in the native *pha* locus (gM1). High activity of the native enzymes within module M2 (gM2) was facilitated by promoter engineering of the corresponding genes (*PP_0989*-*PP_0986*). Elements in the outline are not drawn to scale; gene deletions are indicated with a red cross. *Km^R^*, kanamycin-resistance determinant; *RBS*, ribosome binding site; *BCD10*, bicistronic design used as a translational coupler. **(c)** A synthetic C1-glycine-serine auxotrophy is rescued upon expression of modules M1 and M2 with formate as the only source of glycine. Engineered *P*. *putida* strains were cultivated in 96-well microtiter plates in de Bont minimal medium with the supplements indicated in each case [10% (v/v) CO_2_ in the headspace, 20 mM glucose, 2 mM glycine and/or 30 mM formate]. The prefix ‘p’ or ‘g’ indicates expression of the corresponding modules either from a plasmid or the chromosome, respectively. Average values for bacterial growth (estimated as the optical density measured at 600 nm, OD_600_) and doubling times (DT) ± standard deviation of three biological replicates are represented. **(d)** Growth profile of the engineered strains in shaken-flask cultures is directly coupled to both **(e)** glucose and **(f)** and **(g)** formate consumption. Cultivations were performed in biological triplicates in de Bont minimal medium supplemented with 20 mM glucose and 30 mM formate in a 10% (v/v) CO_2_ atmosphere. Average values ± standard deviation for the specific consumption rate of glucose (*q*_G_) and formate (*q*_F_), as well as DTs, are indicated in the figure. *CDW*, cell dry weight; *concn*., concentration.

The combined activity of the first and second modules (M1 and M2) of the synthetic rGly pathway should also relieve the auxotrophy in *P*. *putida* SLA and enable growth in glucose cultures upon addition of formate as the sole glycine source (**Fig. 3a**). Note that all experiments involving this assimilation route require cultivation of the engineered strains in a CO_2_-enriched atmosphere, as module M2 of the rGly pathway incorporates an extra C1 unit in the form of CO_2_ (**Fig. 2**). The low-copy-number plasmid pSFM1 (**Table 1**), carrying module M1 as a synthetic operon under the control of the constitutive P_4_ promoter [a derivative of P*_trc_* (57)], was implemented to explore this possibility (**Fig. 3b**). Introducing plasmid pSFM1 in strain SLA, however, could not restore growth of *P*. *putida* SLA in DBM with glucose and formate (**Fig. 3c**). This result prompted the question of whether the GCS activity (which forms the core of the M2 module in the assimilation pathway) could be a bottleneck towards C1 incorporation.

2-Oxoacid dehydrogenases (e.g. the pyruvate dehydrogenase complex, the 2-oxoglutarate dehydrogenase complex and the branched-chain α-ketoacid dehydrogenase complex) typically convert 2-oxoacid substrates to the corresponding acyl-coenzyme A (CoA) form and yield NADH and CO_2_. The GCS belongs to this category of dehydrogenases and, in the context of formate assimilation, the GCS activity should run in the reverse, anabolic direction (i.e. glycine → pyruvate). Owing to their key role in catabolic pathways, genes encoding this type of 2-oxoacid dehydrogenases are subject to stringent transcriptional control (58). There are two annotated GCSs in the genome of *P*. *putida* KT2440 (59), and the genes encoding GcvH (protein H), GcvP (glycine dehydrogenase) and GcvT (aminomethyltransferase) are located in two distant loci in the chromosome. Cluster I (*PP_0986*-*PP_0989*) further contains a gene encoding L-serine dehydratase (*tdcG-II*), whereas cluster II (*PP_5192*-*PP_5194*) is solely formed by *gcvP-II*, *gcvH-II* and *gcvT-II*. Although the transcriptional regulation of these genes remain unexplored, we found that *PP_0997*, encoding a σ^54^ dependent transcriptional regulator/sensory box protein, lies close to cluster I. Sequence analysis revealed that the PP_0997 protein shares a high degree of homology (79%) with the GcvR regulator of *P*. *aeruginosa* PAO1, known to modulate the expression of genes encoding the (single cluster) GCS in that species (60,61). We adopted two complementary approaches to debottleneck the GCS node in the engineered *P*. *putida* strain: (i) deletion of *PP_0997*, to override any native regulation on either GCS cluster, and (ii) promoter engineering of cluster I by implementing a synthetic P_14g_(*BCD10*) element, to boost the expression of *gcvH-I*, *gcvP-I*, *tdcG-II* and *gcvT-I* (**Fig. 3b**). Furthermore, module M1 was integrated in the bacterial genome (gM1) by a knock-in/knock-out strategy into the native *pha* locus (*PP_5002*-*PP_5008*; **Fig. 3b** and **Fig. S2** in the Supplemental Data). This chromosomal manipulation fulfills two purposes. On the one hand, the *pha* locus is an adequate genome spot for stable gene integration and expression from the chromosome. On the other hand, deleting the *pha* gene cluster blocks polyhydroxyalkanoate (PHA) accumulation—which would otherwise act as a carbon sink, competing with biomass formation (62).

The combination of these engineering efforts led to *P*. *putida* strain gM1gM2 [*P. putida* SLA Δ*PP_0997* P_14g_(*BCD10*)→*gcv-I* Δ*PP_5002-PP_5008*::P_4_→*mtdA*-*fch*-*ftfL* from *M*. *extorquens*; **Table 1**]. Expression of the two modules from the chromosome, combined with supplementation of 30 mM formate as the sole source of C1 moieties, glycine and serine, restored growth of *P*. *putida* strain gM1gM2 cultured in DBM medium containing 20 mM glucose and 10% (v/v) of CO_2_ in microtiter plate cultures (**Fig. 3c**). Interestingly, expressing M1 as a genome-integrated, single copy (gM1) did not alter growth kinetics (DT = 2.4 ± 0.1 h) in comparison to expression from the low-copy-number plasmid pSFM1 (DT = 2.3 ± 0.1 h). Likewise, the C1 assimilation module provided enough glycine for biomass build-up as did direct supplementation of the amino acid, since the final cell densities and growth patterns were essentially identical in both cases (**Fig. 3c**). This engineered strain could not grow in any of the control conditions tested (i.e. glucose and formate in the absence of CO_2_, or glucose or formate in a CO_2_-enriched atmosphere; **Fig. 3c**), indicating that glucose is still needed to supply both energy and part of the biomass precursors— with the C1 feedstock replenishing the glycine pool and metabolites downstream.

Further characterization of growth parameters and substrate consumption was carried out in shaken-flask cultures using the same cultivation conditions described above (i.e. glucose and formate in a CO_2_-enriched incubator). The characteristic DTs of *P*. *putida* pM1gM2 and gM1gM2 were very similar to each other (**Fig. 3d**), albeit slightly longer than those observed in microtiter plate cultures. The strain engineered with modules M1 and M2 as chromosomal insertions had a 2.3-fold faster specific glucose consumption rate (*q*_G_) than the strain carrying plasmid pSFM1 (**Fig. 3e**), qualitatively mirroring formate consumption patterns (i.e. *q*_F_ was the highest in *P*. *putida* gM1gM2; **Fig. 3f** and **Fig. 3g**). With this physiological characterization at hand, we set to explore whether formate is actually incorporated into biomass (rather than oxidized) by adopting ^13^C-tracer experiments.

### 3.3. ^13^C-labeled C1 tracers underscore the origin and fate of C atoms in the metabolic network

In order to dissect the contribution of the synthetic modules to biomass formation in the engineered *P*. *putida* strains, the distribution of formate in proteinogenic amino acids was assessed by feeding ^13^C-labeled C1 tracers—either as formate or CO_2_ (**Fig. 4a**). In particular, the pattern of ^13^C enrichment in glycine, serine and alanine was analyzed by LC-MS after biomass hydrolysis. As expected, after cultivating the engineered strains in the presence of ^12^C-glucose, ^13^C-formate and ^12^CO_2_, glycine is labeled only once—whereas serine and part of alanine are labeled twice (**Fig. 4b**). As observed in the growth experiments described above (Fig. 3), the labeling patterns did not differ between strains pM1gM2 and gM1gM2—and, in both cases, 100% of the glycine pool was directly derived from formate assimilation. Cultivation in the presence of ^12^C-glucose, ^12^C-formate and ^13^CO_2_ led to similar labeling configurations except for alanine, which was labeled once, indicating a significant flux entering central metabolism *via* serine deamination to pyruvate (**Fig. 4c**). Upon cultivation with ^12^C-glucose, ^13^C-formate and ^13^CO_2_, glycine and serine were predominantly labeled twice and three times, respectively, while ca. 15% of the alanine pool was labeled three times (**Fig. 4d**). As expected, the wild-type strain is naturally unable to assimilate formate, leading to no labeling upon ^13^C-formate supplementation. The pool of glycine and serine was, however, partially labeled when wild-type cells were incubated in a ^13^CO_2_-enriched atmosphere (ca. 25-40%), likely due to the anaplerotic flux forming oxaloacetate from pyruvate and CO_2_ that leads to threonine, which can be cleaved off to yield glycine. Taken together, these ^13^C-labelling results confirm biosynthesis of glycine, serine and alanine from the synthetic formate assimilation module when glucose as the main carbon source.

**Figure 4.**
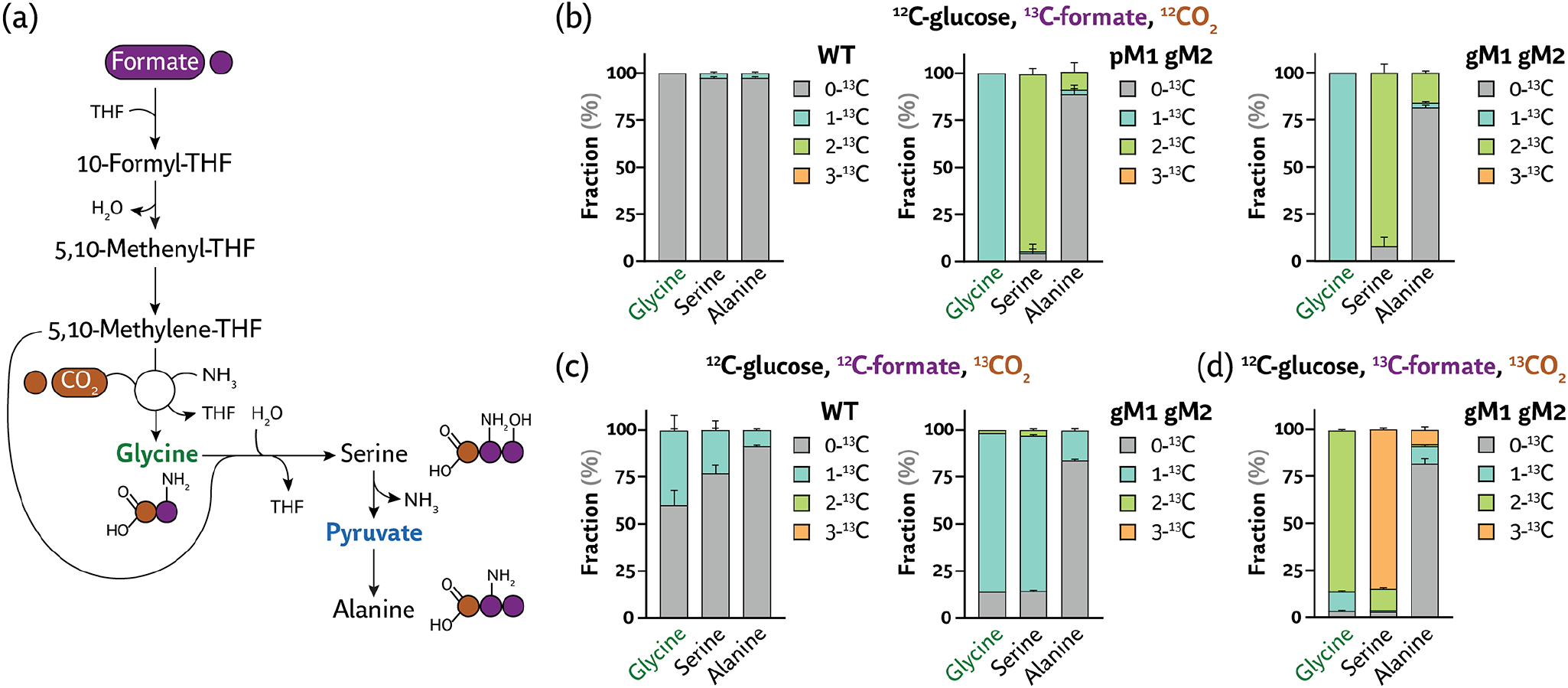
^13^C-tracer experiments reveal the pattern of formate assimilation into glycine, serine and alanine. **(a)** Metabolic origin of glycine, serine and alanine, the key proteinogenic amino acids stemming from the synthetic rGly pathway. ^13^C-labelled C1 tracers (^13^C-formate or ^13^CO_2_) fed to engineered *P*. *putida* strains, expressing synthetic module M1 (either on a plasmid format, pM1, or from the genome, gM1) and module 2 (gM2), were used to identify the origin and fate of C atoms in the biochemical network by analyzing the pattern of ^13^C enrichment in the three amino acids indicated. *THF*, tetrahydrofolate. **(b)** Labeling pattern in wild-type (strain EM42, WT) and engineered *P*. *putida* strains cultivated in the presence of ^13^C-formate and unlabeled glucose and CO_2_. Cells were incubated in 24-well deep-well plate cultures and biomass samples were collected upon cultures reached stationary phase. Samples were hydrolyzed in 6 M HCl and the pattern of ^13^C enrichment (indicated as a percentage of the total pool) was analyzed by LC-MS. **(c)** Labeling pattern upon cultivating either the wild-type (strain EM42, WT) or the engineered *P*. *putida* strain in an atmosphere enriched in ^13^CO_2_. **(d)** Labeling pattern upon cultivating the *P*. *putida* strains under study in the presence of ^13^CO_2_ and ^13^C-formate. In all cases, average values for the fraction of ^13^C enrichment ± standard deviation of three biological replicates are presented.

As further of the synthetic pathway functionality, ribose was tested as an alternative substrate to provide carbon and energy instead of hexoses. To this end, the engineered *P*. *putida* strains were grown under the same conditions described previously, but with 24 mM ribose replacing glucose (but providing an equimolar amount of C atoms). The DTs of strains pM1gM2 and gM1gM2 increased ca. 4-fold as compared to those observed in glucose cultures (**Fig. S3a** in the Supplemental Data). Since ribose is slowly assimilated by *P*. *putida*, high incorporation of ^13^C-formate into pyruvate was observed in these experiments—as indicated by the fact that 25% of the alanine pool was labeled twice (**Fig. S3b**). These results further support the validity of the modular approach adopted towards synthetic formatotrophy in *P*. *putida*, and indicate that the fraction of C1 assimilated through the synthetic rGly route also depends on the nature of the primary carbon source. On this basis, our next objective was to increase the fraction of biomass derived from formate by introducing further manipulations in the engineered strains.

### 3.4. Establishing formate as the main carbon source in engineered P. putida

Encouraged by the significant carbon flux *via* the rGly route to pyruvate, we aimed at increasing the selection strength on the pathway. By omitting glucose, all carbon needs to be generated *via* the synthetic rGly pathway. To slightly lower the metabolic demand, we supplied acetate to the medium in addition to formate; since the glyoxylate shunt was blocked (Δ*aceA* deletion, **Table 1**), acetate can only contribute to a limiting number of biomass precursors—i.e. acetyl-CoA and 2-ketoglutarate. Thus, the strategy relies on increasing the demand of C units derived from the synthetic formatotrophy modules such that no secondary carbon source is needed for biosynthesis of biomass precursors—while a surrogate substrate is used as an energy source. In these conditions, the C2 feedstock can only be oxidized in the TCA cycle to produce reducing equivalents and energy while the C1 substrate supports biomass formation (**Fig. 5a**). A combined gene deletion-and-overexpression strategy was implemented to this end, as explained below.

**Figure 5.**
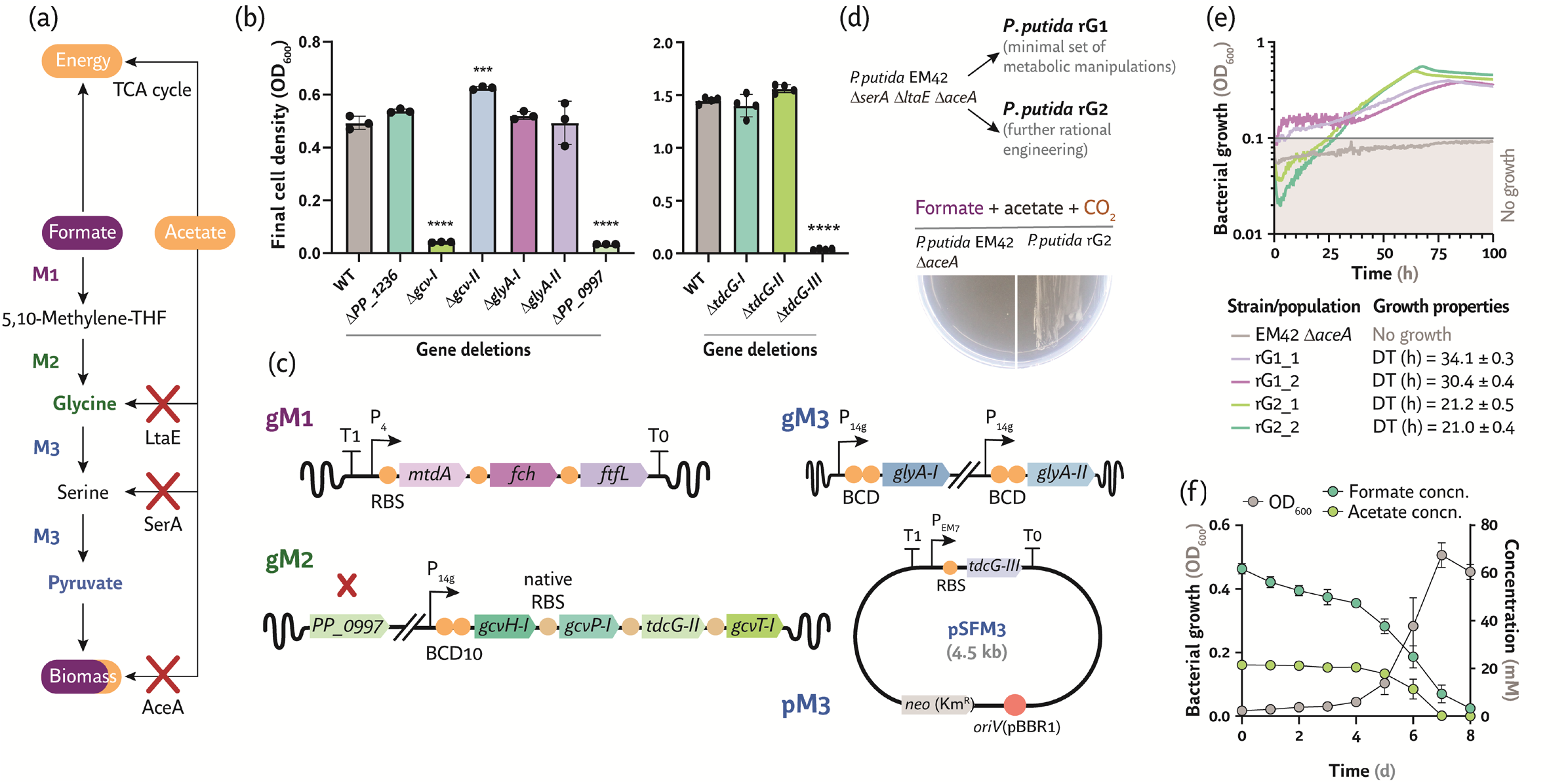
Engineering formate assimilation in *Pseudomonas putida via* the synthetic rGly pathway. **(a)** Tight selection strategy to assimilate formate as main carbon source by deleting *ltaE*, *serA* and *aceA* (*P*. *putida* SLA) and overexpressing modules M1, M2 and M3 in different configurations (genome-based or plasmid-borne). **(b)** Role of the (redundant) glycine cleavage systems, GCS regulators, serine hydroxymethyl transferase and L-serine deaminases in the growth of *P*. *putida* on amino acids. The final cell density (at 24 h, estimated as the optical density at 600 nm, OD_600_) of *P. putida* EM42 (WT) harboring deletions in *gcv-I*, *gcv-II*, *PP_1236*, *PP_0997*, *glyA-I*, *glyA-II* and each of the *tdcG* genes is represented for three biological replicates ± standard deviation with ∗ *p* < 0.05, ∗∗ *p* < 0.01, ∗∗∗ *p* < 0.001 and ∗∗∗∗ *p* < 0.0001. Each strain was cultivated in 96-well microtiter plates in DBM medium supplemented with 20 mM glycine (*gcv*, *glyA* and GCS regulator mutants) or 20 mM serine (*tdcG* mutants). **(c)** First formatotrophic design implemented in *P*. *putida*. Modules M1, M2, *glyA-I* and *glyA-II* were constitutively expressed from the genome and *tdcG-III* from a medium-copy-number, broad-host-range plasmid (pSFM3, **Table 1**). Additional deletions (*PP_0997*) are indicated with a red cross. *Km^R^*, kanamycin-resistance determinant; *RBS*, ribosome binding site; *BCD*, bicistronic design used as a translational coupler. **(d)** Two lineages of synthetic *P*. *putida* formatotrophs were constructed on the background of strain SLA. In strains rG1 and rG2, *glyA-I* and *glyA-II* are expressed from different bicistronic designs (*BCD2* and *BCD20*, respectively; **Table 1**). The full description of the lineages is provided in **Fig. S2** in the Supplemental Data. Growth of a synthetic glycine auxotroph of *P*. *putida* (strain rG1) is rescued upon expression of modules M1, M2 and M3 in the presence of formate as the only source of biomass precursors. The *aceA* deletion prevents assimilation of acetate, which acts mostly as energy source. The strains under study (*P*. *putida* rG2 and Δ*aceA*, used as a control) were streaked in solid de Bont minimal medium with 30 mM formate and 20 mM acetate. Plates were incubated for 2 weeks at 30°C in a 10% (v/v) CO_2_ atmosphere and photographed afterwards. **(e)** Growth profile of engineered *P*. *putida* strains (either rG1 or rG2; **Table 1**) upon adaptive laboratory evolution (ALE) in the presence of formate and acetate. Evolved populations were cultivated in 96-well microtiter plates in de Bont minimal medium supplemented with 30 mM formate and 20 mM acetate and 10% (v/v) CO_2_ in the headspace. Average values for bacterial growth (estimated as the OD_600_) and doubling times (DT) ± standard deviation of three replicates are represented. rG1_1 and rG1_2, and rG2_1 and rG2_2, refer to two bacterial populations evolved in parallel. **(f)** Shaken-flask cultures of (minimally) engineered and evolved *P*. *putida* incubated under formatotrophic conditions. Evolved clone rG1_1/2 was incubated in de Bont minimal medium supplemented with 60 mM formate and 20 mM acetate in a 10% (v/v) CO_2_ atmosphere. Cultivations were performed in biological triplicates, and average values ± standard deviation for OD_600_ and substrate concentration (concn.) are indicated in the figure.

#### 3.4.1. Dissecting the role of native metabolic functions in C1 assimilation

As indicated previously, the core of the rGly pathway is formed by the enzymes of the GCS running in the carboxylating direction. While experiments in **Fig. 3** indicate that overexpressing genes that encode the GCS-I ensures enough flux through this pathway to provide all C1, glycine and serine, we were interested in gaining insight on the role of the *gcv-I* and *gcv-II* clusters of *P*. *putida*. Thus, we deleted individually the genes that encode the two GCSs, the annotated (*PP_1236*) and the putative (*PP_0997*) GCS regulators of strain EM42 (**Fig. 5b**). Eliminating GCS-I or *PP_0997* abolished growth of the mutants in DBM medium added with 20 mM glycine, whereas deletion of genes encoding GCS-II or *PP_1236* had no effect on the phenotype. As such, at this stage we focused on *gcv-I* overexpression and deletion of *PP_0997*. Module M3 is also required to ensure enough flux from glycine to pyruvate. Module M3 encodes two enzymes, GlyA (L-serine hydroxymethyltransferase) and TdcG (L-serine deaminase). Genes annotated to encode these activities in strain KT2440 are *glyA-I* (*PP_0322*), *glyA-II* (*PP_0671*), *tdcG-I* (*PP_0297*), *tdcG-II* (*PP_0987*) and *tdcG-III* (*PP_3144*). As the deletion of each L-serine hydroxymethyltransferase individually didn’t affect the phenotype of the mutant on DBM medium supplemented with 20 mM serine (**Fig. 5b**), and the transcriptional regulation patterns for these genes is yet to be explored, we decided to overexpress the two genes encoding L-serine hydroxymethyltransferase *via* promoter engineering. To this end, a strong, constitutive promoter (P_14g_) and a translational coupler (*BCD2* or *BCD20*) were integrated upstream of *glyA-I* and *glyA-II* in *P*. *putida* gM1gM2, leading to strain rG1 [*P. putida* SLA Δ*PP_0997* P_14g_(*BCD10*)→*gcv-I* Δ*PP_5002-PP_5008*::P_4_→*mtdA*-*fch*-*ftfL* from *M*. *extorquens* and P_14g_*(BCD2)*→*glyA-I P_14g_(BCD2)*→*glyA-II*; **Table 1**]. In parallel, we attempted to identify which of the three genes encoding TdcG should be overexpressed to ensure proper conversion of serine into pyruvate by individually deleting each of the three *tdcG* genes of *P*. *putida* EM42 (i.e. the parental strain selected for engineering formatotrophy). The resulting strains were grown in DBM medium with 20 mM serine. Deletion of *tdcG-I* and *tdcG-II* did not hamper growth of the strains as they reached a final cell density comparable to *P*. *putida* EM42 strain (**Fig. 5b**), whereas the Δ*tdcG-III* strain was unable to grow even after prolonged (>72 h) incubation. These results suggest that TdcG-III is essential for growth on serine, and we overexpressed *tdcG-III* to enhance pyruvate supply from the synthetic rGly pathway. Plasmid pSFM3, a medium-copy-number vector carrying a P_EM7_→*tdcG-III* element (**Table 1** and **Fig. 5c**), was implemented to this end.

#### 3.4.2. Rational manipulation of the carbon and redox balances towards formate assimilation

*P*. *putida* rG1 was able to grow on DBM medium plates supplemented with 30 mM formate and 20 mM acetate and incubated in a 10% (v/v) CO_2_ atmosphere, with colonies clearly discernable after ca. 2 weeks of cultivation (**Fig. 5d**). Expectedly, *P*. *putida* EM42 Δ*aceA* (**Fig. 5d**) or strain SLA could not grow under these culture conditions. These results indicate that establishing synthetic C1 assimilation in *P*. *putida* depends on multiple metabolic factors, and we decided to explore the minimal set of modifications that could enable formate conversion into biomass. To this end, two lineages of formatotrophic *P*. *putida* strains were constructed (**Fig. 5d**). On the one hand, *P*. *putida* rG1 was selected as a minimally-engineered lineage while, on the other hand, *P*. *putida* rG2 was obtained by modifying the rG1 background with five targeted manipulations. These modifications, targeting not only carbon assimilation but also redox balance, are: (i) overproduction of L-serine deaminase by means of the P_EM7_→*tdcG-III* element encoded in plasmid pSFM3, (ii) overexpression of the second gene cluster encoding the GCS *via* chromosomal promoter engineering [i.e. implementing P_14g_*(BCD10)*→*gcv-II*, to further boost the carboxylation step carried by the GCSs], (iii) deletion of genes encoding the regulators of central carbon metabolism Crc (*PP_5292*, catabolite repression control protein) and HexR (*PP_1021*), (iv) elimination of potential competing pathways for formate consumption by deleting *purT* [*PP_1457*, encodes a formate-dependent (5-phospho-β-D-ribosyl)glycinamide formyltransferase] and (iv) enhancing NADPH turnover *via* transhydrogenation by means chromosomal promoter engineering [i.e. implementing P_EM7_→*pntAA* (*PP_0156*)]. In this design, both Crc and HexR were eliminated to avoid potential bottlenecks in glycolytic fluxes, such that pyruvate from the synthetic rGly pathway can feed the upper metabolism *via* gluconeogenesis in the EDEMP cycle (63), the central metabolic architecture of *P*. *putida*. Likewise, the misrouting of the C1 feedstock into the formation of *N*^2^-formyl-*N*^1^-(5-phospho-β-D-ribosyl)glycinamide was abolished (Δ*purT*), while the electron transfer from NADH to NADP^+^ generating NADPH, the cofactor required for methylene-THF biosynthesis, was fostered by constitutive expression of the membrane-bound, NADPH-producing pyridine nucleotide transhydrogenase. All these metabolic modifications, as well as the genealogy of the formatotrophic strains constructed in this study, is summarized in **Fig. S2** in the Supplemental Data. Next, we focused on integrating an evolution approach to further boost C1 assimilation in engineered *P*. *putida* strains.

### 3.5. Adaptive laboratory evolution enables efficient formate assimilation in two lineages of engineered P. putida

The rG1 and rG2 lineages are designed to decouple C1 assimilation from energy formation (the latter being supported by acetate feeding), and both strains grew very slowly on formate as the main carbon source. In order to reduce DTs of the engineered strains, *P*. *putida* rG1 and rG2 were submitted to adaptive laboratory evolution (ALE) by repeated passage of the cultures in DBM medium supplemented with 30 mM formate and 20 mM acetate with 10% (v/v) CO_2_ in the headspace. After the ALE experiment (lasting ca. 3 months), the evolved bacterial populations were regrown under the same conditions to check stability of the growth phenotypes (**Fig. 5e**). Individually-evolved populations are identified with an underscore (i.e. rG1_1 and rG1_2 represent two parallel evolution experiments from strain *P*. *putida* rG1; rG2_1 and rG2_2 are evolved populations derived from *P*. *putida* rG2). ALE significantly accelerated the growth rates (thereby reducing DTs) of both engineered strains under formatotrophic conditions, and different populations evolved from the same lineage displayed similar DTs and growth patterns. All of the bacterial populations analyzed reached a similar final cell density (OD_600_ ≈ 0.3-0.5). However, we observed that the rG1_1 and rG1_2 evolved populations (derived from the *P*. *putida* strain with minimal metabolic interventions) had a DT ≈ 30 h, whereas the rG2_1 and rG2_2 populations (obtained from the heavily engineered strain rG2) had the shortest DT ≈ 20 h (**Fig. 5e**). *P*. *putida* Δ*aceA* was included as a negative control to account for the small variations in OD_600_ readings that could occur in prolonged cultivation in microtiter plate readers.

Next, we aimed at identifying specific mutations that could lead to these phenotypic differences. To this end, nine independent clones were isolated and purified from evolved populations of each of the engineered strains and their growth on formate, acetate and CO_2_ was compared under the same conditions indicated in previous experiments. Importantly, all 18 clones were passaged on complex LB medium before assessing formatotrophic growth, such that the observed phenotypes could be attributed to mutations accumulated during evolution and not to transient adaptation phenomena. Clones isolated from the rG1_1 and rG1_2 populations exhibited heterogeneity in their formatotrophic growth patterns (**Fig. S4a** in the Supplemental Data), whereas all individual *P*. *putida* colonies purified from the evolved rG2 populations recapitulated the growth profile observed when growing the entire population (**Fig. S4b** in the Supplemental Data). Whole-genome sequencing of individual clones was performed upon extraction of chromosomal DNA, and mutations were annotated and catalogued to identify differences between the parental strain and the corresponding evolved progeny. All relevant mutations identified in evolved clones are listed in **Fig. S5** in the Supplemental Data. The heterogeneity in growth patterns of rG1 clones was reflected in the landscape of mutations therein. A substitution in *folM* (*PP_4632*), encoding a bifunctional dihydrofolate/7,8-dihydromonapterin reductase that catalyzes the dihydrofolate to tetrahydrofolate conversion required for condensation with formate, was identified in clone rG1_1/2 (note that, in this strain nomenclature, the last number indicates the individual clone under consideration). The second alteration is a deletion in *sdhA* (*PP_4191*), the flavoprotein subunit of the succinate dehydrogenase complex in the TCA cycle. Other clones of the same population accumulated other mutations that could potentially exert a pleiotropic effect. In clone rG1_1/5, for instance, we detected a single substitution in *PP_0264*, a sensor histidine kinase of unknown function. As similar sensor histidine kinase systems are involved in amino acid or dicarboxylic acids uptake, we hypothesized that this mutation could be involved in the uptake of key metabolites of the rGly pathway. As indicated above, the nine clones isolated from the rG2_1 and rG2_2 populations had similar growth pattern and identical mutations were identified by whole-genome sequencing. Pleiotropic modifications were detected in *glnK* (*PP_5234*), encoding the NR^II^(GlnL/NtrB) phosphatase activator involved in nitrogen uptake and regulation, *PP_t01*, corresponding to a tyrosine-specific tRNA, and in *PP_2166*, an anti-σ factor antagonist. A substitution in *glnE* (*PP_0340*), encoding glutamate/NH_4_^+^-ligase adenylyltransferase and involved in ammonium assimilation, was also identified in clone rG2_1/2. While these mutations could certainly modify the formatotrophic phenotypes of evolved clones by fine-tuning carbon and redox metabolism, we focused on unraveling the main catabolic pathways involved in glycine processing as explained in the next section.

### 3.6. Parallel evolution trajectories support formate assimilation in engineered P. putida via glycine hydroxymethylation or oxidation

We wanted to further confirm the phenotype of rG1_1/2 observed from the microtiter plate experiments in shaken-flask cultures using formate as the carbon substrate and acetate as the energy source. Cultures of *P*. *putida* rG1_1/2 followed the trend previously observed for the entire evolved population, with full consumption of both formate and acetate after 1 week of incubation under formatotrophic conditions (**Fig. 5f**)—concomitant with the highest cell density observed thus far (OD_600_ ≈ 0.6). Using labelled tracers, we decided to study the pattern of ^13^C assimilation in the parallel-evolved isolates rG1_1/2 and rG2_2/5 to analyze if (i) acetate (expected to act mostly as energy source, owing to the Δ*aceA* deletion in the engineered strains) could be partially assimilated and/or (ii) glycine could be channeled into alternative catabolic routes other than the expected conversion into pyruvate. The biochemical fate of carbon atoms from the C1 and C2 feedstock—specifically, in serine, alanine, proline, asparagine and histidine—was explored by ^13^C-tracer experiments upon feeding cultures of all these isolates with ^12^CO_2,_ ^12^C/^13^C-formate and/or ^12^C/1-^13^C-acetate. 1-^13^C-acetate was supplemented to simplify the analysis as the carbonyl carbon of labeled acetate remains in the first TCA cycle turn but is completely lost in the second turn whereas the methyl carbon remains during two full turns, before losing one half in each succeeding turn of the cycle. Under these conditions, intermediates within the upper metabolism should be entirely derived from formate, whereas the TCA cycle metabolites could partially reflect the ^13^C-signal from acetate if there is assimilation of this feedstock (**Fig. 6a**). We likewise contemplated the possibility that glycine oxidation, rather than its conversion into pyruvate *via* hydroxymethylation followed by deamination by GlyA and TdcG, respectively, could emerge as an assimilation bypass in the absence of AceA (**Fig. 6a**). The genome of *P*. *putida* encodes a FAD-dependent glycine/D-amino acid oxidase, ThiO (*PP_0612*), which could potentially catalyze the oxidative conversion of glycine into glyoxylate.

**Figure 6.**
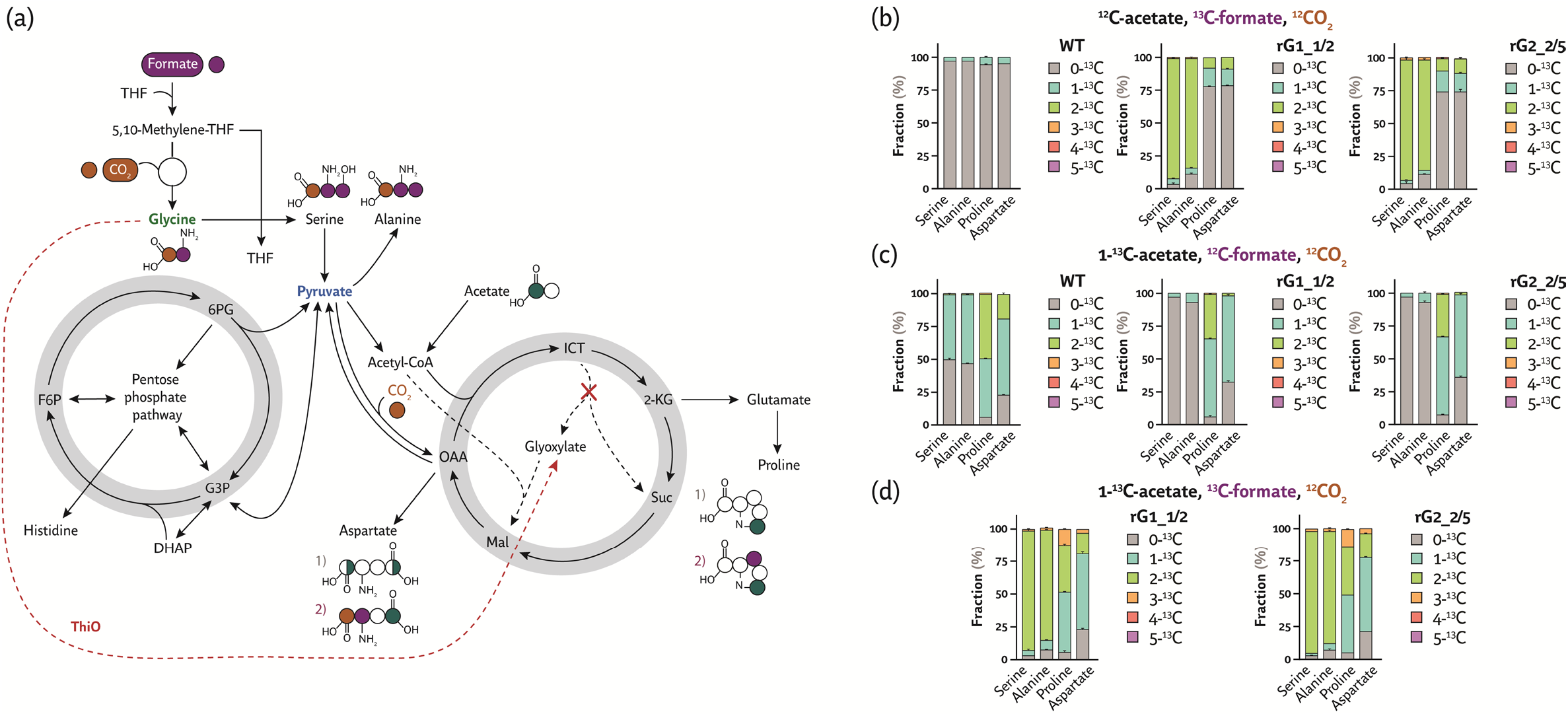
Pyruvate is mainly derived from the C1 feedstock in engineered *Pseudomonas putida* grown under formatotrophic conditions. **(a)** Proteinogenic amino acids derived from the synthetic rGly pathway and the core metabolism of *P*. *putida*. The metabolic origin of the amino acids shown in the scheme can be tracked by ^13^C-tracer experiments in the engineered strain expressing modules M1, M2 and M3 (evolved clones rG1_1/2 and rG2_2/5) upon feeding ^12^CO_2_, ^12^C/^13^C-formate and/or ^12^C/1-^13^C-acetate. Note that a different labeling pattern of proline and aspartate is expected depending on whether ThiO is inactive (*1A*) or active (*2*). **(b)** Labeling pattern in wild-type (WT) and engineered *P*. *putida* strains (isolated rG1_1/2 and rG2_2/5 clones) fed with ^13^C-formate and unlabeled acetate. Cells were incubated in 24-deep-well plate cultures and biomass samples were collected upon reaching early stationary phase. Samples were hydrolyzed in 6 M HCl and the pattern of ^13^C enrichment (indicated as a percentage of the total pool) was analyzed by LC-MS. **(c)** Labeling pattern upon cultivating the *P*. *putida* strains under study in the presence of 1-^13^C-acetate and unlabeled formate. **(d)** Labeling pattern upon cultivating the engineered *P*. *putida* strains in the presence of ^13^C-formate and 1-^13^C-acetate. In all cases, average values for the fraction of ^13^C enrichment ± standard deviation of three biological replicates are presented.

During cultivation of the engineered strains in the presence of ^13^C-formate, ^12^C-acetate and ^12^CO_2_, serine (∼91%), alanine (∼83%) and part of proline (∼8%) and aspartate (∼10%) pools were labeled twice (**Fig. 6b**). For the wild-type strain (EM42), however, ∼5% of proline, ∼5% of aspartate and ∼3% of alanine were labeled only once, most likely corresponding to natural distribution of ^13^C-carbons in acetate. In the presence of ^12^C-formate, 1-^13^C-acetate and ^12^CO_2_, proline was either labeled once or twice while aspartate was labeled only once. Feeding the cultures with ^13^C-formate and 1-^13^C-acetate led to a labeling profile of serine and alanine that was similar to that observed with labeled formate. The ^13^C-content in proline, aspartate and histidine, in contrast, was increased due to potential incorporation of both labeled formate and acetate (**Fig. 6d**). These results indicate that serine and alanine are derived from formate, while asparagine and proline seem to be mainly derived from acetate. Interestingly, the labeling patterns obtained for clones rG1_1/2 and rG2_2/5 were essentially identical. The role of ThiO, if any, is difficult to directly derive from these experiments due to, among other factors, the cycling of the ^13^C signal through the pyruvate shunt and the EDEMP cycle (63,64).

To investigate the potential branching of catabolism between the expected glycine conversion into serine and pyruvate or ThiO-dependent oxidation (**Fig. 7a**), we adopted a genetic approach to dissect the fate of the amino acid. Deleting *thiO* in strain EM42 did not affect glycine assimilation (**Fig. 7b**), indicating that ThiO does not play a major catabolic role in non-evolved *P*. *putida*. Next, *thiO* was individually deleted in clones rG1_1/2 and rG2_2/5, and the growth profile of the resulting strains was compared in DBM medium supplemented with formate, acetate and CO_2_ (**Fig. 7c**). Deletion of *thiO* in *P*. *putida* rG1_1/2 increased the DT by 2-fold and reduced the cell density by 17% under formatotrophic conditions. Eliminating ThiO in clone rG2_2/5, increased the DT from 26.5 ± 0.1 h to 80.1 ± 0.1 h (i.e. 3-fold longer). These results suggest the notion that glycine assimilation *via* GlyA and TcdG prevails in the rG1 lineage, whereas glycine oxidation *via* ThiO is the main catabolic fate of the amino acid in evolved clones of strain rG2. Taken together, these observations indicate that different catabolic modes for C1 assimilation emerged in *P*. *putida* depending on the metabolic background of the strains before ALE optimization of formatotrophic growth.

**Figure 7.**
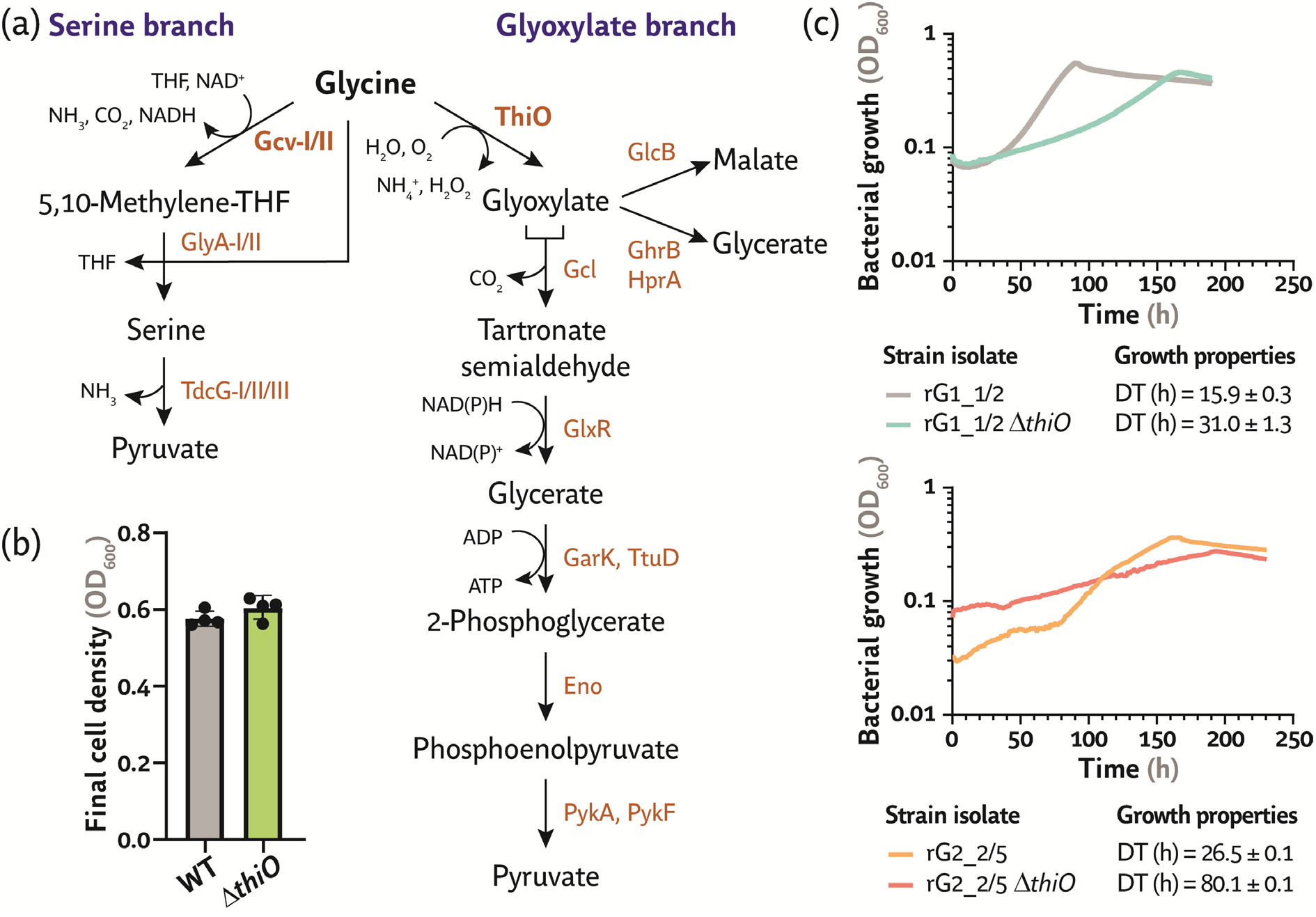
C1 assimilation through the glycine cleavage system and ThiO in evolved clones of engineered *P*. *putida*. **(a)** Glycine can be metabolized by *P. putida via* the glycine cleavage system (either Gcv-I and Gcv-II) or oxidized by ThiO (annotated as a FAD-dependent glycine/D-amino acid oxidase). Either serine or glyoxylate can be produced, depending on which branch of glycine processing is active. **(b)** Final cell density (estimated as the optical density at 600 nm, OD_600_) of *P. putida* EM42 (WT) and its D*thiO* derivative. Cells were cultivated for 24 h in 96-well microtiter plates in DBM medium supplemented with 20 mM glycine, and the average cell density of three biological replicates ± standard deviation is represented together with individual data points. **(c)** Growth phenotypes of evolved clones isolated from two formatotrophic *P*. *putida* lineages and their D*thiO* derivatives. *P*. *putida* rG1_1/2, rG1_1/2 D*thiO*, rG2_2/5 and rG2_2/5 D*thiO* were cultivated in DBM medium supplemented with 60 mM formate and 20 mM acetate in a 10% (v/v) CO_2_ atmosphere in 96-well microtiter plates. Average values for bacterial growth (OD_600_) and doubling times (DT) ± standard deviation of three biological replicates are shown.

## 4. Conclusion

There is no doubt that valorizing CO_2_ by biotechnological processes is the way forward to tackle the global challenge of greenhouse gas emission (6,22). Electrochemical conversion of CO_2_ into soluble C1 molecules followed by assimilation by microorganisms lies at the core of this strategy (15). In this study, we expanded the substrate repertoire of reduced-genome *P*. *putida* EM42 to include formate—the most promising soluble C1 feedstock owing to its solubility in water and efficient production from CO_2_ (65). Over time, *P*. *putida* has been established as a robust host for bioproduction due to its capability to thrive in a variety of environmental conditions, a feature matched by high levels of resistance to oxidative stress and toxic chemicals (30). *P. putida* is also characterized by a versatile metabolism allowing for the assimilation of a broad variety of substrates that range from simple sugars and amino acids to arene molecules (66)—however, C1 feedstock is not part of this extensive substrate repertoire. Integrating rational engineering, together with implementing synthetic auxotrophies relieved upon functional expression of metabolic modules (67) and short-term laboratory evolution (68), enabled functional decoupling of C1 assimilation from energy generation.

We initially divided the rGly pathway into three metabolic modules and confirmed the activity of each segment by relieving synthetic auxotrophies established by gene deletion(s). Incorporation of formate and CO_2_ into biomass in the presence of glucose was confirmed by ^13^C-tracer experiments and analysis of the pattern of ^13^C enrichment in amino acids. In this first design, formate-derived glycine was entirely processed through the action of the GCS encoded by *gcv-I*. Based on these results, we constructed two lineages of engineered strains that contained either a minimal set of metabolic manipulations (rG1) or several modifications introduced to optimize carbon and redox metabolism (rG2). The DT of the second generation of engineered *P*. *putida* formatotrophs cultivated on formate as the main carbon source and acetate for energy generation was reduced to ca. 15 h by combining rational engineering with ALE. Multiple mutations that could lead to the improved growth phenotype were identified, yet the number and nature of DNA modifications that consistently appeared across isolates from evolved populations were relatively small. Due to the function of structural and regulatory proteins affected by these mutations (e.g. PP_0264), together with altered activity of enzymes that likewise accumulated modifications during evolution (e.g. SdhA), these adaptations could exert a global effect on carbon fluxes.

Regardless of the presence of such adaptive mutations, we found that the rG1 and rG2 lineages evolved to rely on two different catabolic regimes at the C1-derived glycine node. Upon evolution, *P*. *putida* rG1 processed glycine partly through its conversion to serine and pyruvate (i.e. the intended route), whereas rG2 clones mainly adapted to transform the amino acid into glyoxylate (i.e. by-passing the Δ*aceA* modification in this strain that was expected to fully block acetate incorporation, as indicated by the patterns of ^13^C incorporation). As such, ThiO acted as an alternative pathway in evolved *P*. *putida* rG1 while being the main catabolic route for glycine assimilation in strain rG2. The theoretical study by Cotton et al. (15) indicated that the activity of the glyoxylate branch of glycine processing would result in a decrease of 11% of the biomass yield, 22% of the pyruvate yield and 25% of the acetyl-CoA yield as compared to the serine branch. Further investigation would be needed to fully unravel the reasons underlying such an interesting evolutionary feature. The glyoxylate branch seems to be the less preferred option for glycine processing—not only is this route composed by several metabolic steps to yield pyruvate (seven, as compared to only three in the serine branch), but it also generates reactive oxygen species in the form of H_2_O_2_ while consuming NADPH. Notably, this pathway predominantly emerged in the rG2 lineage, which was heavily modified at the level of carbon and redox metabolism. Taken together, these experimental observations indicate that the metabolic background adopted for engineering formatotrophy dictates which catabolic route(s) will ultimately lead to biomass generation upon ALE. As such, the results of this article illustrate the portability of the rGly pathway while highlighting the impact of the metabolic background where synthetic formatotrophy is to be engineered. These results will pave the way to establish C1 assimilation in non-model microbial platforms, adding to the wealth of cell factories that could make the formate bioeconomy a tangible reality in the near future.

## Supplemental Data

**Table S1.** Oligonucleotides used in this study

**Table S2.** Codon-optimized gene sequences used for engineering *P*. *putida*

**Fig. S1 ·** Glycine and serine supplementation rescues synthetic auxotrophies in *P*. *putida*

**Fig. S2 ·** Genealogy of synthetic formatotrophic *P*. *putida* strains

**Fig. S3 ·** Growth of the selection *P*. *putida* strains on formate, ribose and CO_2_

**Fig. S4 ·** Adaptive laboratory evolution of a *P*. *putida* strain engineered for growth on formate, acetate and CO_2_

**Fig. S5 ·** Whole genome sequencing analysis of evolved *P*. *putida* clones engineered for formate assimilation

## Authors’ Contributions

J.T. and P.I.N. conceived the project and designed the experiments. J.T., B.D. and A.D.M. performed the experiments leading to the results described in the article. All the authors analyzed the data and participated in the discussions included in this study. J.T. and P.I.N. wrote the manuscript, with contributions of all the other authors.

## Acknowledgments

We thank Nicolas T. Wirth, Mariela P. Mezzina and Fiorella Masotti for their help with the construction of some of the plasmids reported in this work. We also thank Änne Michaelis for her help in LC-MS analysis of amino acids. The financial support from The Novo Nordisk Foundation through grants NNF20CC0035580, *LiFe* (NNF18OC0034818) and *TARGET* (NNF21OC0067996), the Danish Council for Independent Research (*SWEET*, DFF-Research Project 8021-00039B), and the European Union’s Horizon 2020 Research and Innovation Programme under grant agreement No. 814418 (*SinFonia*) to P.I.N. is gratefully acknowledged. J.T. is the recipient of a fellowship from The Novo Nordisk Foundation as part of the Copenhagen Bioscience Ph.D. Programme, supported through grant NNF17CC0026768.

## Footnotes

1 We refer to ‘formate’ in the text for the sake of consistency, as sodium formate was used as the main carbon source, yet the actual form assimilated *via* the synthetic pathway is formic acid.

