## Supplemental Data 1 for "Integrated rational and evolutionary engineering of genome-reduced *Pseudomonas putida* strains empowers synthetic formate assimilation"

by

Justine Turlin[a], Beau Dronsella[b], Alberto De Maria[a], Steffen N. Lindner[c], and Pablo I. Nikel[a]*

[a] The Novo Nordisk Foundation Center for Biosustainability, Technical University of Denmark, Kongens Lyngby, Denmark
[b] Max Planck Institute for Terrestrial Microbiology, Marburg, Germany
[c] Charité Universitätsmedizin, Berlin, Germany

**Table S1.** Oligonucleotides used in this study.

| ***Name*** | **DNA sequence** (5'→3') | **Use** |
|---|---|---|
| *serA*-U-US-F | AGATCCUTGGCAAACCACAGTACGGT | Construction of pGNW2·Δ*serA* |
| *serA*-U-US-R | ACCTTACAUCTGCGTAAACCTGTATCCC | |
| *serA*-U-DS-F | ATGTAAGGUCGCTGGCGGTATGAAAAAGG | |
| *serA*-U-DS-R | AGGTCGACUGCCGTTGCCGGTCTGCTGAT | |
| *ltaE*-U-US-F | AGATCCUAGCCTTCGGCGTACTTGT | Construction of pGNW2·Δ*ltaE* |
| *ltaE*-U-US-R | ATGGCACGGUCCTGTGAACGACGCAGAT | |
| *ltaE*-U-DS-F | ACCGTGCCAUGTGAGCGAGAGCCGG | |
| *ltaE*-U-DS-R | AGGTCGACUGCTCGCGTTGAACGCT | |
| *aceA*-U-US-F | AGATCCUTGTCGACTGATCCCAGGCAAT | Construction of pGNW2·Δ*aceA* |
| *aceA*-U-US-R | ATCACAUGGAAATAAACCTCGTCGCATCGA | |
| *aceA*-U-DS-F | ATGTGAUAGTGGGTTGCCGTTGAAC | |
| *aceA*-U-DS-R | AGGTCGACUCAGGATGTCACCGAGTACCGAGAA | |
| *PP_1236*-U-US-F | AGATCCUCGGTGCCGGCGATGACCGGAAT | Construction of pGNW2·Δ*PP_1236* |
| *PP_1236*-U-DS-F | ATGCAGCAUTCCTAAAGCGATGAGCGGTCGGC | |
| *PP_1236*-U-DS-R | ATGCTGCAUGTAAGGAAACCGATCATGG | |
| *PP_1236*-U-DS-R | AGGTCGACUCAGCCCCATTCAATTTCA | |
| *PP_0997*-U-US-R | ACATGGGGUTCGAACAGTTGGCAATGG | Construction of pGNW2·Δ*PP_0997* |
| *PP_0997*-U-DS-F | ACCCCATGUAACGACTGTGGTCAAAACCCATAA | |
| *PP_0997*-U-DS-R | AGGTCGACUCCATCAGCGGGTCGAGCC | |
| *PP_0997*-U-DS-R | AGGTCGACUCCATCAGCGGGTCGAGCC | |
| *crc*-U-US-F | ACCTCCAGUGATGATCTGCATGACCTCACGAATGG | Construction of pGNW2·Δ*crc* |
| *crc*-U-US-R | ACATAAAUGGCCCCATAAATCTCGTG | |
| *crc*-U-DS-F | ATTTATGUAAAAGGCCATTGGGGCTGCAT | |
| *crc*-U-DS-R | AGCTCTTGAUAACGCCATGCTCGCTTTGGC | |
| *purT*-US-U-F | AGATCCUCTACGCCTTCTCCAGTGCC | Construction of pGNW2·Δ*purT* |
| *purT*-US-U-R | ATTCAGGUCCTCGAAGGCATCCGGGG | |
| *purT*-DS-U-F | ACCTGAAUCCGAGAGTGCTGGGGCCG | |
| *purT*-DS-U-R | AGGTCGACUCCGCGAAACGTTCCTTGGCC | |
| *gcvI*-U-US-F | AGATCCUTGTTCCCGCTATTGCTGGTGCTCG | Construction of pGNW2·Δ*gcv-I* |
| *gcvI*-U-US-R | ATGGGGAAUCCTCGGGAAGCAGGGCGT | |
| *gcvI*-U-DS-F | ATTCCCCAUGTGAAACGGCTTTGTGAAG | |
| *gcvI*-U-DS-R | AGGTCGACUTCAGACAGGTTGAACCTT | |
| *gcvII*-U-US-F | AGATCCUGCGCTTCATCGCGGTGGC | Construction of pGNW2·Δ*gcv-II* |
| *gcvII*-U-US-R | AACAGGGUTTCTCCTTCCGGGCGTGGC | |
| *gcvII*-U-DS-F | ATCGTTTCUAATGGGACAGCGCACGCTTTTGTAT | |
| *gcvII*-U-DS-R | AGGTCGACUATGAGTGCAGCGCGCGCG | |

| | | |
|---|---|---|
| *glyAI*-U-US-F | AGATCCUACACCTCGCAGTCGGTTTCGAACA | Construction of pGNW2·$\Delta$*glyA-I* |
| *glyAI*-U-US-R | ACTCACATUGTGTATCTCCCGGCAGCGATCGT | |
| *glyAI*-U-DS-F | AATGTGAGUGGAGTCACACACCATGCA | |
| *glyAI*-U-DS-R | AGGTCGACUCAGTCCAGGTACGGGATT | |
| *glyAII*-U-US-F | AGATCCUAGATCGACCGGACCTTCGT | Construction of pGNW2·$\Delta$*glyA-II* |
| *glyAII*-U-US-R | ATCACAUGGCGGTCCTCAAGGATCG | |
| *glyAII*-U-DS-F | ATGTGAUTGGCGCTTGCCTAAAGTGA | |
| *glyAII*-U-DS-R | AGGTCGACUTCAGGCGTGAGTCGCCTTT | |
| $P_{EM7}$_*sdaIII*-F | AGTATAATACGACAAAAGCTTAGGAGGAAAAACAT ATGGCTATCAGTGTTTTCGACCTTT | Insertion of the $P_{EM7}$ promoter in plasmid pSFM3 |
| $P_{EM7}$_*sdaIII*-R | ATGCCGATATACTATGCCGATGATTAATTGTCAAC ATTAATTAAAGGCATCAAATAAAAC | |
| *gcvII*-US-U-F | AGATCCUGCGCTTCATCGCGGTGGC | Construction of pGNW2·*$P_{14g}$ (BCD10)→ gcv-II* |
| *gcvII*-US-U-R | AACAGGGUTTCTCCTTCCGGGCGTGGC | |
| *gcvII*-DS-U-F | ATCGTTTCUAATGGGACAGCGCACGCTTTTGTAT | |
| *gcvII*-DS-U-R | AGGTCGACUATGAGTGCAGCGCGCGCG | |
| $P_{EM7}$-BCD10-U-F | ACCCTGTUGACAATTAATCATCGGCATA | |
| $P_{EM7}$-BCD10-U-R | AGAAACGAUCCTCCGCATGATTAAGAT | |
| *tdcG-I*_U_US-F | AGATCCUTGGCGCGGACAGCAACGC | Construction of pGNW2·$\Delta$*tdcG-I* |
| *tdcG-I*_U_US-R | ATCACATUGTCGAATCACCTCTTGCTGGGCT | |
| *tdcG-I*_U_DS-F | AATGTGAUTTTCTGAGCTGACCTTCT | |
| *tdcG-I*_U_DS-R | AGGTCGACUGCGATTTCTCCTCACACA | |
| *tdcG-II*_U_US-F | AGATCCUGACCATGTCCTTCCCGGT | Construction of pGNW2·$\Delta$*tdcG-II* |
| *tdcG-II*_U_US-R | ATGTGGACCUCCTGCCAGGTTGTTGGGT | |
| *tdcG-II*_U_DS-F | AGGTCCACAUGTGACCCGACAATGACAGCAAGGA | |
| *tdcG-II*_U_DS-R | AGGTCGACUACCTTGACTGCAGCCGGG | |
| *tdcG-III*_U_US-F | AGATCCUAAAACGCGGCAAGCGCGG | Construction of pGNW2·$\Delta$*tdcG-III* |
| *tdcG-III*_U_US-R | ATCACAUGCTAATGCCTGACCCTGCCCT | |
| *tdcG-III*_U_DS-F | ATGTGAUGTTTAACGCGGTCAATTGGAGC | |
| *tdcG-III*_U_DS-R | AGGTCGACUTGCAGGAAATCACCCGCTACT | |
| P4-*ftfL*_ins_*pha*-F | TGACATCAGGAAAATTTTTCTGTATAATGTGTGGA | Construction of pGNW2·$\Delta$*pha*::M1 |
| P4-*ftfL*_ins_*pha*-R | CTAGAACAGCCCGTCGATCTGACC | |
| P14g-BCD2-U-R | AGAAAACCUCCTTAGCATGATTAAGATGTTTCA | Construction of pGNW2·*$P_{14g}$(BCD2) →glyA-I* |
| *glyAI*_U-R | AGGTCGACUTTGTGCTCGACCGCCAGG | |
| *glyAI*_US_U-F | AGATCCUAACCGGACGGTGCAGGGGATT | |
| *glyAI*_US_U-R | ATGGGCTGUGTATCTCCCGGCAGCGATCGT | |
| *glyAI*_P14gBC_U-F | ACAGCCCAUTGACAAGGCTCTCGCGGC | |
| *glyAI*_U_F | AGGTTTTCUAATGTTCAGCAAGCAAGACCAGAT | |

| | | |
|---|---|---|
| P14g-BCD2-U-R | AGAAAAACCUCCTTAGCATGATTAAGATGTTTCA | Construction of pGNW2· *$P_{14g}$(BCD2)* →*glyA-II* |
| *glyAII*_US_U-F | AGATCCUAGATCGACCGGACCTTCGT | |
| *glyAII*_U-F | AGGTTTTCUAATGTTCAGCCGTGATTTGACCAT | |
| *glyAII*_U-R | AGGTCGACUGGCTTGTGCTCGACAGCC | |
| *glyAII*_US_U-R | ATGGGCGGCGGUCCTCAAGGATCGGGGTAATTTG | |
| *glyAII*_PBC_U-F | ACCGCCGCCCAUTGACAAGGCTCTCGCGGC | |
| P14gBC_*glyAI*-F | GAAAGTTTCTAATGTTCAGCAAGCAAGACCAGAT | Construction of pGNW2· *$P_{14g}$(BCD20)* →*glyA-I* |
| P14gBC_*glyAI*-R | TCAGCATGATTAAGATGTTTCAGTACGAAAATTGC | |
| P14gBC_*glyAII*_F | GAAAGTTTCTAATGTTCAGCCGTGATTTGACC | Construction of pGNW2· *$P_{14g}$(BCD20)* →*glyA-II* |
| P14gBC_*glyAII*_R | TCAGCATGATTAAGATGTTTCAGTACGAAAATTGC | |
| pGNW2-PBG-F | CGGCATAGTATATCGGCATAGTATAATACGACAAGGGCCCAAGTTCACTTAAAAAGG | Construction of pGNW2· *$P_{14g}$ (BCD10)*→ *gcv-I* |
| pGNW2-PBG-R | ATGATTAATTGTCAACAGGGGAATCCTCGGGAAGCAGGGCGTT | |
| $P_{EM7}$_*pntAB*_F | CGATATACTATGCCGATGATTAATTGTCAACAGACTATTTCTCCTGCGGTGACCTTTTG | Construction of pGNW2· *$P_{EM7}$-pntAA* |
| $P_{EM7}$_*pntAB*_R | GCATAGTATAATACGACAAAAGCTTAGGAGGAAAAACATGTGCACATTGGTGTTCCTCTC | |
| *pha*_U_US-F | AGATCCUCCTGCAGTTCGGCAAGATCAACGT | Construction of pGNW2· Δ*pha* |
| *pha*_U_US-R | ATGCATCUACGACGCTCCGTTGTCCTGAGA | |
| *pha*_U_DS-F | AGATGCAUGCTGTGTACCTCATGCTCAT | |
| *pha*_U_DS-R | AGGTCGACUAACACATGGGGTGGGCTGAT | |
| *thiO*_U_US-F | AGATCCUGGTTAGCCTGTTCCGGATTAC | Construction of pGNW2· Δ*thiO* |
| *thiO*_U_US-R | AGGTCCUCACATCTGTCGATCCAACACCTGAAG | |
| *thiO*_U_DS-F | AGGACCUCTTCGCGGGTAAACCCGCTC | |
| *thiO*_U_DS-R | AGGTCGACUGCAGAAAGCGTTGCTTGACCC | |
| pGNW-USER-F | AGTCGACCUGCAGGCATGCAAGCTTCT | Amplification of vector pGNW2 |
| pGNW-USER-R | AGGATCUAGAGGATCCCCGGGTACCG | |

**Table S2.** Codon-optimized gene sequences used for engineering *P. putida*.

| **Enzyme** | **UniProt entry** | **DNA sequence** (5'→3') |
|---|---|---|
| Formate-tetrahydrofolate ligase from *Methylobacterium extorquens* AM1 | Q83WS0 | ATGCCCTCAGATATCGAGATCGCCCGCGCGGCGACCCTGA<br>AGCCGATCGCCCAGGTCGCCGAAAAGCTCGGCATCCCGGA<br>CGAGGCGCTTCACAACTACGGCAAGCACATCGCCAAGATC<br>GACCACGACTTCATCGCCTCGCTCGAGGGTAAGCCCGAGG<br>GCAAGCTGGTGCTCGTCACCGCGATCTCGCCGACGCCCGC<br>GGGCGAGGGCAAGACCACCACGACCGTGGGTCTCGGCGAC<br>GCACTCAACCGGATCGGCAAGCGGGCGGTGATGTGCCTGC<br>GCGAGCCCTCGCTCGGCCCCTGCTTCGGCATGAAGGGCGG<br>CGCGGCCGGTGGCGGCAAGGCCCAGGTCGTGCCGATGGAG<br>CAGATCAACCTGCACTTCACCGGGGACTTCCACGCCATCA<br>CCTCGGCGCACTCGCTCGCCGCCGCGCTGATCGACAACCA<br>CATCTACTGGGCCAACGAGCTCAACATCGACGTGCGCCGC<br>ATCCACTGGCGCCGCGTGGTCGACATGAACGACCGGGCGC<br>TGCGCGCGATCAACCAGTCGCTCGGCGGCGTCGCCAACGG<br>CTTTCCGCGTGAGGACGGGTTCGACATCACCGTCGCCTCC<br>GAGGTGATGGCGGTGTTCTGCCTCGCCAAGAATCTGGCCG<br>ACCTCGAGGAGCGGCTCGGCCGCATCGTCATCGCCGAGAC<br>CCGCGACCGCAAGCCGGTGACGCTGGCCGACGTGAAGGCG<br>ACCGGCGCGATGACCGTTCTCCTCAAGGATGCGCTGCAGC<br>CGAACCTCGTGCAGACGCTGGAGGGCAACCCGGCCCTGAT<br>CCATGGCGGCCCGTTCGCCAACATCGCCCACGGCTGCAAC<br>TCGGTGATCGCCACCCGTACCGGCCTGCGGCTGGCCGACT<br>ACACCGTCACCGAGGCCGGCTTCGGCGCGGATCTCGGCGC<br>GGAGAAGTTCATCGACATCAAGTGCCGCCAGACCGGCCTC<br>AAGCCCTCGGCGGTGGTGATCGTCGCCACGATCCGCGCCC<br>TCAAGATGCATGGCGGCGTCAACAAGAAGGATCTCCAGGC<br>TGAGAACCTCGACGCGCTGGAGAAGGGTTTCGCCAACCTC<br>GAGCGCCACGTGAACAACGTGCGGAGCTTCGGCCTGCCGG<br>TGGTGGTGGGCGTGAACCACTTCTTCCAGGACACCGACGC<br>CGAGCATGCCCGGTTGAAGGAGCTCTGCCGCGACCGTCTT<br>CAGGTCGAGGCGATCACCTGCAAGCACTGGGCGGAGGGCG<br>GCGCGGGCGCCGAGGCTCTGGCGCAGGCCGTGGTGAAGCT<br>CGCCGAGGGCGAGCAGAAGCCGCTGACCTTCGCCTACGAA<br>ACTGAGACGAAGATCACCGACAAGATCAAGGCGATCGCGA<br>CCAAGCTCTACGGTGCGGCCGATATCCAGATCGAGTCGAA<br>GGCCGCCACCAAGCTCGCCGGCTTCGAGAAGGATGGCTAC<br>GGCGGATTGCCCGTCTGCATGGCCAAGACGCAGTACTCGT<br>TCTCGACCGACCCGACCCTGATGGGCGCGCCCTCGGGCCA<br>CCTCGTCTCGGTGCGCGACGTGCGCCTCTCGGCGGGCGCC<br>GGCTTCGTCGTGGTGATCTGCGGTGAGATCATGACCATGC<br>CGGGCCTGCCCAAGGTGCCGGCGGCGGACACCATCCGCCT<br>CGACGCCAACGGTCAGATCGACGGGCTGTTCTAG |

| | | |
|---|---|---|
| 5,10-Methenyltetrahydrofolate cyclohydrolase from *Methylobacterium extorquens* AM1 | P55818 | ATGGCCGGCAACGAGACGATCGAAACATTCCTCGATGGCC<br>TGGCGAGCTCGGCCCCGACCCCCGGCGGCGGCGGTGCCGC<br>CGCGATCTCCGGCGCCATGGGCGCGGCGCTGGTCTCGATG<br>GTGTGTAACCTCACCATCGGCAAGAAGAAGTATGTCGAGG<br>TCGAGGCCGACCTGAAGCAGGTGCTGGAGAAGTCGGAAGG<br>CCTGCGCCGCACGCTCACCGGCATGATCGCCGACGACGTC<br>GAGGCTTTCGACGCGGTGATGGGCGCCTACGGGCTGCCGA<br>AAAACACCGACGAGGAGAAGGCTGCCCGCGCCGCCAAGAT<br>TCAGGAGGCGCTCAAGACCGCGACCGACGTGCCGCTCGCC<br>TGCTGCCGCGTCTGCCGCGAGGTGATCGATCTGGCCGAGA<br>TCGTCGCCGAGAAGGGCAATCTCAACGTCATCTCGGATGC<br>CGGCGTCGCCGTGCTCTCGGCCTATGCCGGTCTGCGCTCG<br>GCGGCCCTCAACGTCTACGTCAACGCCAAGGGCCTCGACG<br>ACCGCGCCTTCGCCGAGGAGCGGCTGAAGGAGCTGGAAGG<br>CCTGCTGGCCGAGGCGGGCGCGCTCAACGAGCGGATCTAC<br>GAGACGGTGAAGTCCAAGGTAAACTGA |
| 5,10-Methylenetetrahydrofolate dehydrogenase from *Methylobacterium extorquens* AM1 | P55818 | ATGTCCAAGAAGCTGCTCTTCCAGTTCGACACCGATGCCA<br>CGCCGAGCGTCTTCGACGTCGTCGTCGGCTACGACGGCGG<br>TGCCGACCACATCACCGGCTACGGCAACGTCACGCCCGAC<br>AACGTCGGCGCCTATGTCGACGGCACGATCTACACCCGCG<br>GCGGCAAGGAGAAGCAGTCGACGGCGATCTTCGTCGGCGG<br>CGGCGACATGGCGGCCGGCGAGCGGGTGTTCGAGGCGGTG<br>AAGAAGCGCTTCTTCGGCCCGTTCCGCGTGTCCTGCATGC<br>TGGATTCGAACGGCTCCAACACGACCGCCGCGGCGGGTGT<br>GGCGCTCGTCGTCAAGGCGGCGGGCGGCTCGGTCAAGGGC<br>AAGAAGGCCGTCGTGCTCGCGGGCACCGGCCCGGTCGGCA<br>TGCGCTCGGCGGCGCTGCTCGCCGGCGAGGGCGCCGAGGT<br>CGTGCTGTGCGGGCGCAAGCTCGACAAGGCGCAGGCCGCG<br>GCCGATTCCGTGAACAAGCGCTTCAAGGTGAACGTCACCG<br>CGGCCGAAACCGCGGACGACGCTTCGCGCGCCGAGGCCGT<br>GAAGGGCGCCCATTTCGTCTTCACCGCCGGTGCGATCGGC<br>CTTGAACTGCTGCCGCAGGCAGCCTGGCAGAACGAGAGTT<br>CGATCGAGATCGTGGCCGACTACAACGCCCAGCCGCCGCT<br>CGGCATCGGCGGGATCGATGCGACCGACAAAGGCAAGGAA<br>TACGGCGGAAAGCGCGCCTTCGGTGCGCTCGGCATCGGCG<br>GCTTGAAGCTCAAGCTGCACCGCGCCTGCATCGCCAAGCT<br>GTTCGAGTCGAGCGAAGGCGTCTTCGACGCCGAGGAGATC<br>TACAAGCTGGCCAAGGAAATGGCCTGA |

### Fig. S1 · Glycine and serine supplementation rescues synthetic auxotrophies in *P. putida*

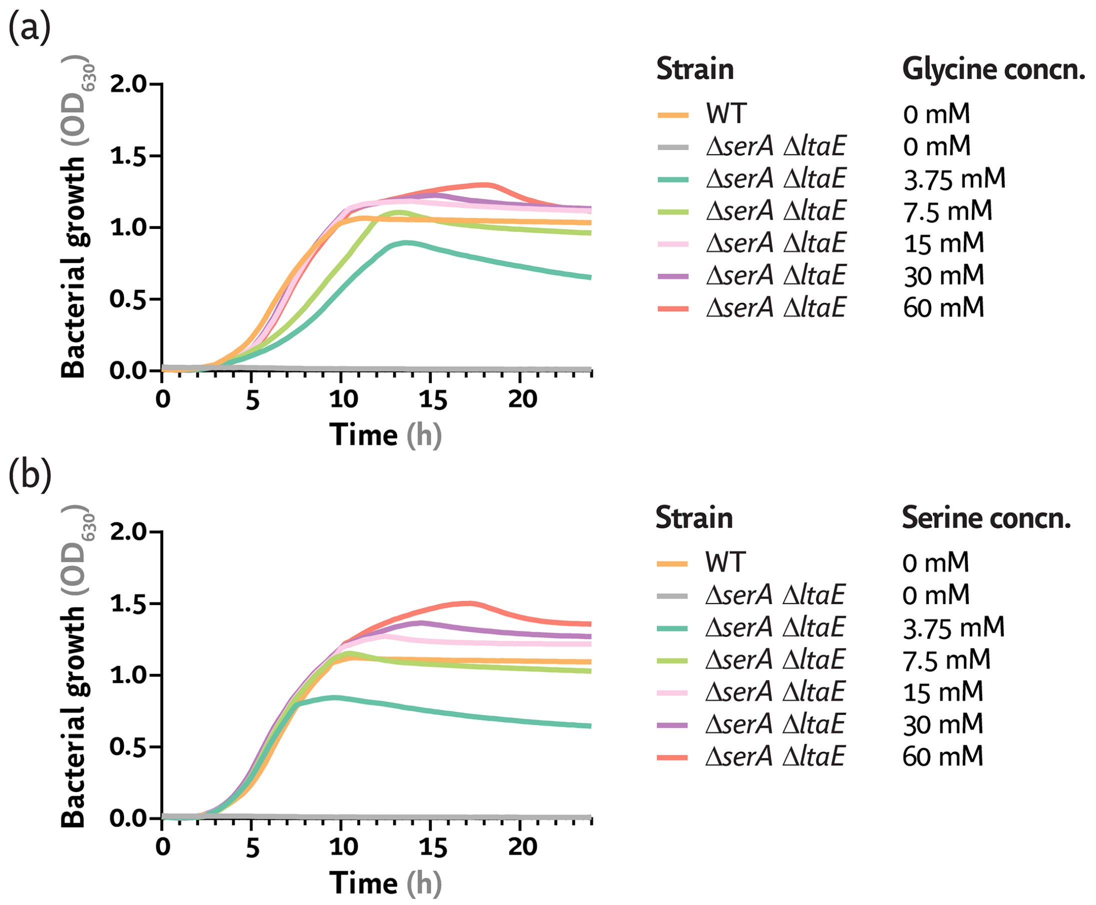


*P. putida* SL is a C1-glycine-serine auxotroph strain (Δ*serA* Δ*ltaE*) derived from genome-reduced *P. putida* EM42. Strain SL is unable to grow in a minimal medium with glucose as the sole carbon source. Supplementation of 3.75, 7.5, 15, 30 or 60 mM of **(a)** glycine or **(b)** serine restores growth of the strain. Wild-type (WT) and engineered *P. putida* strains were cultivated in 96-well microtiter plates in de Bont minimal medium supplemented with 20 mM glucose and glycine or serine when indicated. The average values for bacterial growth (estimated as the optical density measured at 630 nm, $OD_{630}$) of three biological replicates are represented.

### Fig. S2 · Genealogy of synthetic formatotrophic *P. putida* strains

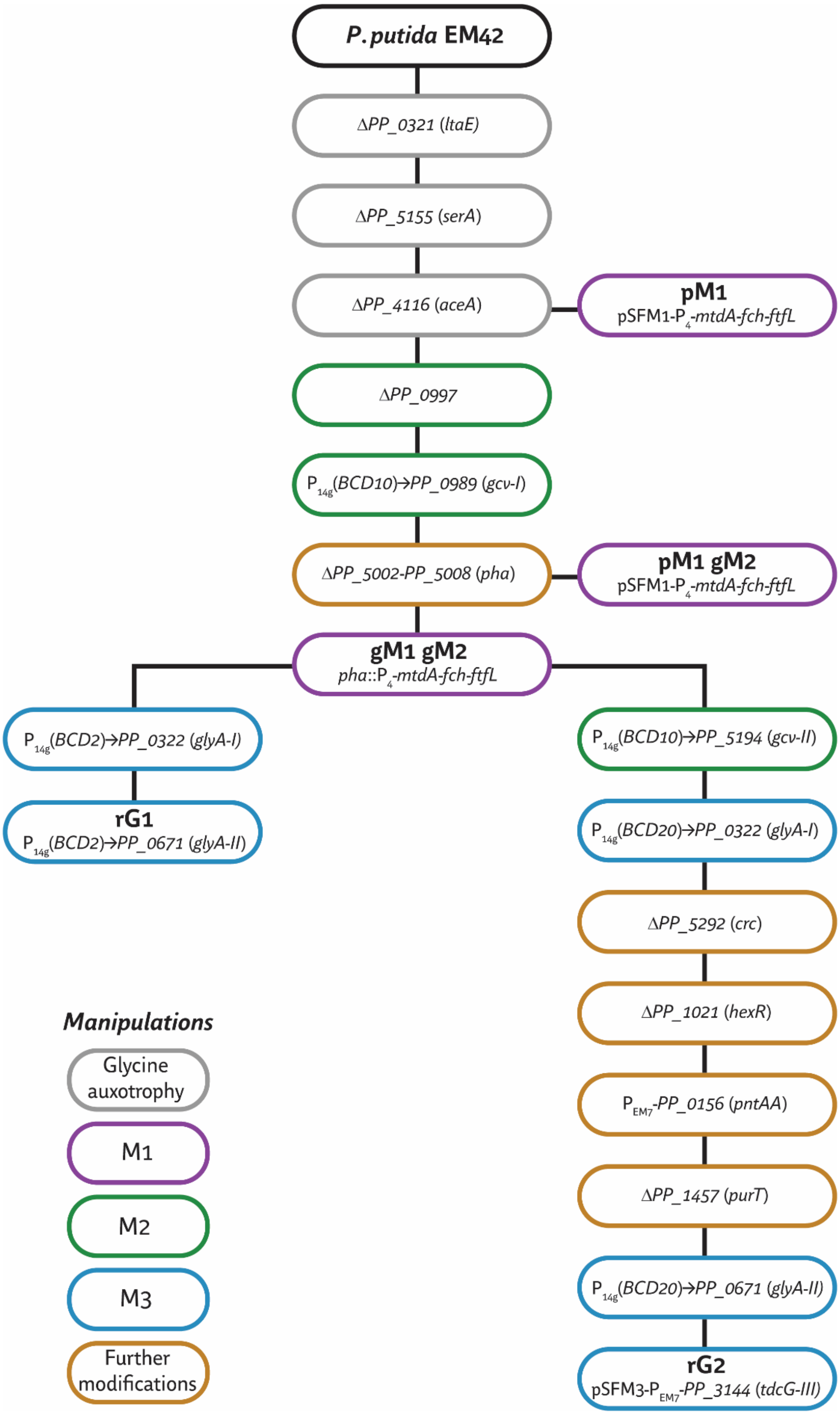


All the strains mentioned in the text and figures of this manuscript are highlighted in bold. Grey circles correspond to deletion(s) required for synthetic glycine auxotrophy. Purple, green and blue circles represent, respectively, modifications involved in modules M1, M2 or M3. Brown circles refer to alterations in global regulation or competing pathways (e.g., redox metabolism).

### Fig. S3 · Growth of the selection *P. putida* strains on formate, ribose and $CO_2$

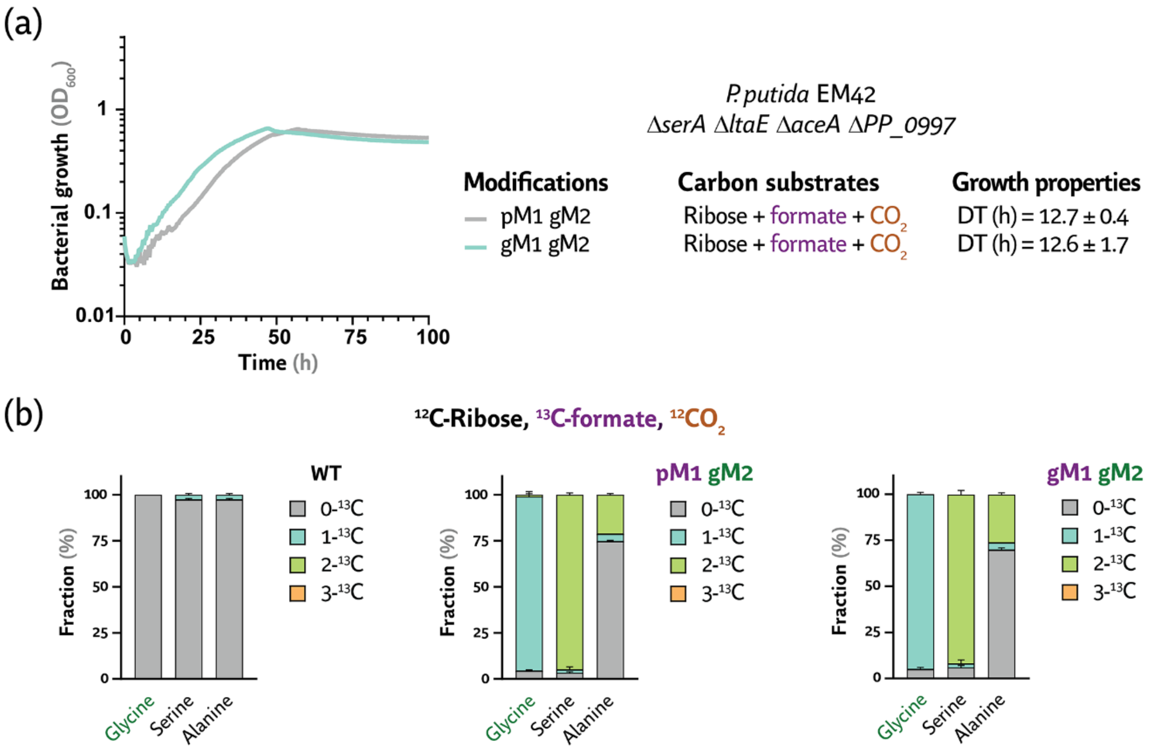


**(a)** Synthetic serine auxotrophy is rescued upon expression of modules M1 and M2 in ribose cultures supplemented with formate (as sole source of glycine) in a $CO_2$-enriched atmosphere. Strains were cultivated in 96-well microtiter plates in de Bont minimal medium with 24 mM ribose, 2 mM glycine and/or 30 mM formate with 10% (v/v) $CO_2$ in the headspace. The average bacterial growth (estimated as the optical density at 600 nm, $OD_{600}$) and average doubling time (DT) ± standard deviation of three biological replicates are represented. **(b)** Labeling pattern in wild-type (WT) and engineered *P. putida* strains cultivated in the presence of $^{13}$C-formate and unlabeled ribose and $CO_2$. Cells were incubated in 24-well deep-well plate cultures, and samples for biomass extraction were collected upon the cultures reached stationary phase. The biomass in these samples was hydrolyzed in 6 M HCl, and the pattern of $^{13}$C enrichment (indicated as a percentage of the total molecule pool) was analyzed by LC-MS.

### Fig. S4 · Adaptive laboratory evolution of a *P. putida* strain engineered for growth on formate, acetate and $CO_2$

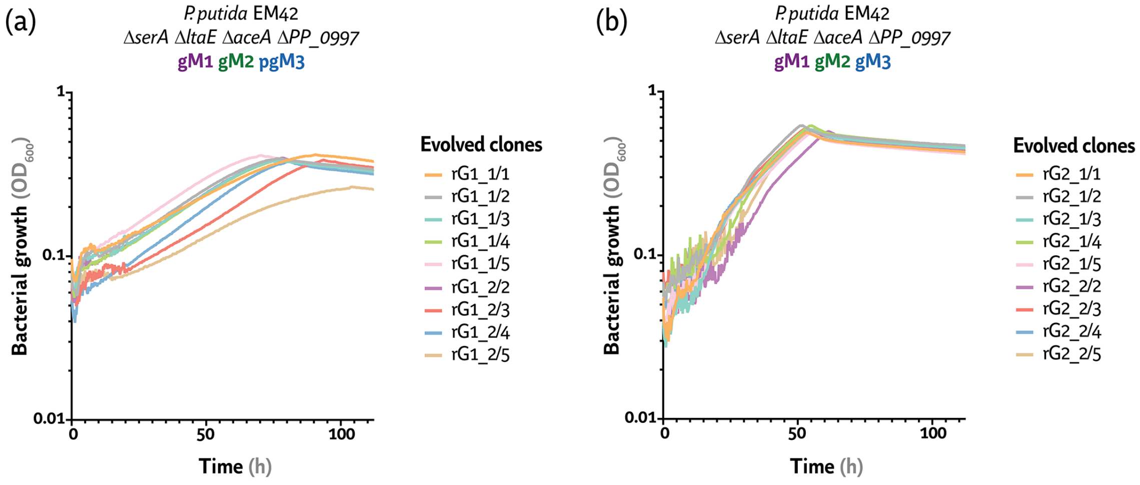


Growth patterns of 18 *P. putida* clones isolated from the evolved populations **(a)** rG1_1 and rG1_2 or **(b)** rG2_1 and rG2_2, cultivated in the presence of formate, acetate and $CO_2$. The evolved clones were incubated in 96-well microtiter plates in de Bont minimal medium supplemented with 60 mM formate and 20 mM acetate with 10% (v/v) $CO_2$ in the headspace. The average bacterial growth (estimated as the optical density at 600 nm, $OD_{600}$) of three experimental replicates are represented.

### Fig. S5 · Whole genome sequencing analysis of evolved *P. putida* clones engineered for formate assimilation

Clone rG1_1/2

| Locus | Function | Sequence | Type |
|---|---|---|---|
| *PP_4632* (*folM*) | Bifunctional dihydrofolate reductase and dihydromonapterin reductase | 5'–GGACTACGCGCCATCATC–3'<br>5'–GGACTACGC**A**CCATCATC–3' | Substitution Ala125Tyr |
| *PP_4191* (*sdhA*) | Succinate dehydrogenase flavoprotein subunit | 5'–CAGGCGGTTCAGGGCAAC–3'<br>5'–CAGGCGG AGGGCAAC–3' | Deletion L56 |

Clone rG1_1/5

| Locus | Function | Sequence | Type |
|---|---|---|---|
| *PP_0264* | Sensor histidine kinase; similar systems are involved in amino acid or dicarboxylate uptake | 5'–CAGGGGCTGGTTCAACTC–3'<br>5'–CAGGGGCTGGT**A**CAACTC–3' | Substitution N→Y |

Clone rG2_1/2

| Locus | Function | Sequence | Type |
|---|---|---|---|
| *PP_0340* (*glnE*) | Glutamate-ammonia ligase adenylyltransferase | 5'–GGCGACCAGGCGGCCGGCGCG–3'<br>5'–GGCG CG–3' | Deletion Δ(AlaTyr)25 DQAAGA→A |
| *PP_5234* (*glnK*) | NRII(GlnL/NtrB) phosphatase activator | 5'–CAGCTTGAACGGCTTGAT–3'<br>5'–CAGCTTGAAC**A**GCTTGAT–3' | Substitution P→L |
| *PP_t01* | tRNA(Tyr) | 5'–TCAAAGGGGACGGACTGT–3'<br>5'–TCAAAGGGG**G**CGGACTGT–3' | Transition |
| *PP_2166* | Anti(anti) σ factor | 5'–TTCGTTGGTGAAGTGCGC–3'<br>5'–TTCGTTGGT**T**AAGTGCGC–3' | Truncation |

Clone rG2_2/5

| Locus | Function | Sequence | Type |
|---|---|---|---|
| *PP_5234* (*glnK*) | NRII(GlnL/NtrB) phosphatase activator | 5'–CAGCTTGAACGGCTTGAT–3'<br>5'–CAGCTTGAAC**A**GCTTGAT–3' | Substitution P→L |
| *PP_t01* | tRNA(Tyr) | 5'–TCAAAGGGGACGGACTGT–3'<br>5'–TCAAAGGGG**G**CGGACTGT–3' | Transition |
| *PP_2166* | Anti(anti) σ factor | 5'–TTCGTTGGTGAAGTGCGC–3'<br>5'–TTCGTTGGT**T**AAGTGCGC–3' | Truncation |

Mutations emerging in clones **(a)** rG1_1/2, **(b)** rG1_1/5, **(c)** rG2_1/2 and **(d)** rG2_2/5 after short-term adaptive laboratory evolution were identified by whole genome sequencing and mapping to the *P. putida* EM42 reference genome. The analysis was performed on three biological replicates in independent isolates from the evolution experiment.
